# A model for the dynamics of expanded CAG repeat alleles: *ATXN2* and *ATXN3* as prototypes

**DOI:** 10.1101/2023.09.07.556735

**Authors:** Lucas Schenatto Sena, Renan Lemes, Gabriel Vasata Furtado, Maria Luiza Saraiva-Pereira, Laura Bannach Jardim

## Abstract

Spinocerebellar ataxia types 2 (SCA2) and 3 (SCA3/MJD) are diseases due to dominant unstable expansions of CAG repeats (CAGexp). Age of onset of symptoms (AO) correlates with the CAGexp length. Repeat instability leads to new expansions, to important AO anticipations and to the eventual extinction of lineages. Because of that, compensatory forces are expected to act on the maintenance of expanded alleles, but they are poorly understood. Here, we described the CAGexp dynamics, adapting a classical equation and aiming to estimate for how many generations will the descendants of a *de novo* expansion last. A mathematical model was adapted to encompass anticipation, fitness, and allelic segregation; and empirical data fed the model. The arbitrated ancestral mutations included in the model had the lowest CAGexp and the highest AO described in the literature. One thousand generations were simulated until the alleles were eliminated, fixed, or 650 generations had passed. All SCA2 lineages were eliminated in a median of 10 generations. In SCA3/MJD lineages, 593 were eliminated in a median of 29 generations. The other ones were eliminated due to anticipation after the 650th generation or remained indefinitely with CAG repeats transitioning between expanded and unexpanded ranges. Therefore, the model predicted outcomes compatible with empirical data - the very old ancestral SCA3/MJD haplotype, and the *de novo* SCA2 expansions -, which previously seemed to be contradictory. This model accommodates these data into understandable dynamics and might be useful for other CAGexp disorders.

## 1. Introduction

CAG repeats expansions (CAGexp) are a major genetic cause of neurological diseases. When they occur within a codon region, the corresponding expansion of the polyglutamine tract (polyQ) in the expressed protein is neurotoxic. Mechanisms include post-transcriptional and -translational modifications and autophagic disturbances (Adegbuyiroa et al 2017; Bunting et al 2021) [1,2]. Each CAGexp or mutant polyQ targets different populations of neurons, causing distinct diseases that are commonly referred to as polyQ diseases. They include Huntington disease (HD, [MIM: 143100]), the spinocerebellar ataxia type 1 (SCA1, [MIM: 164400]), type 2 (SCA2, [MIM: 183090]), type 3 (also known as Machado-Joseph disease, SCA3/MJD, [MIM: 109150]), type 6 (SCA6, [MIM: 183086]), type 7 (SCA7, [MIM: 164500]), type 17 (SCA17, [MIM: 607136]), Dentatorubropallidoluysian atrophy (DRPLA, [MIM: 125370]), and spinobulbar muscular atrophy (SBMA, also known as Kennedy’s disease, [MIM: 313200]) (**Figure 1**) (Orr HT at al 1993; La Spada at al 1991; the Huntington’s Disease Collaborative Research Group. 1993; Kawaguchi at al 1994; Komure et al 1995; David et al 1997; Zhuchenko et al 1997; Koide et al 1999; Pulst et al 2005; Margolis et al 2005) [3,4,5,6,7,8,9,10,11,12].

**Figure 1.**
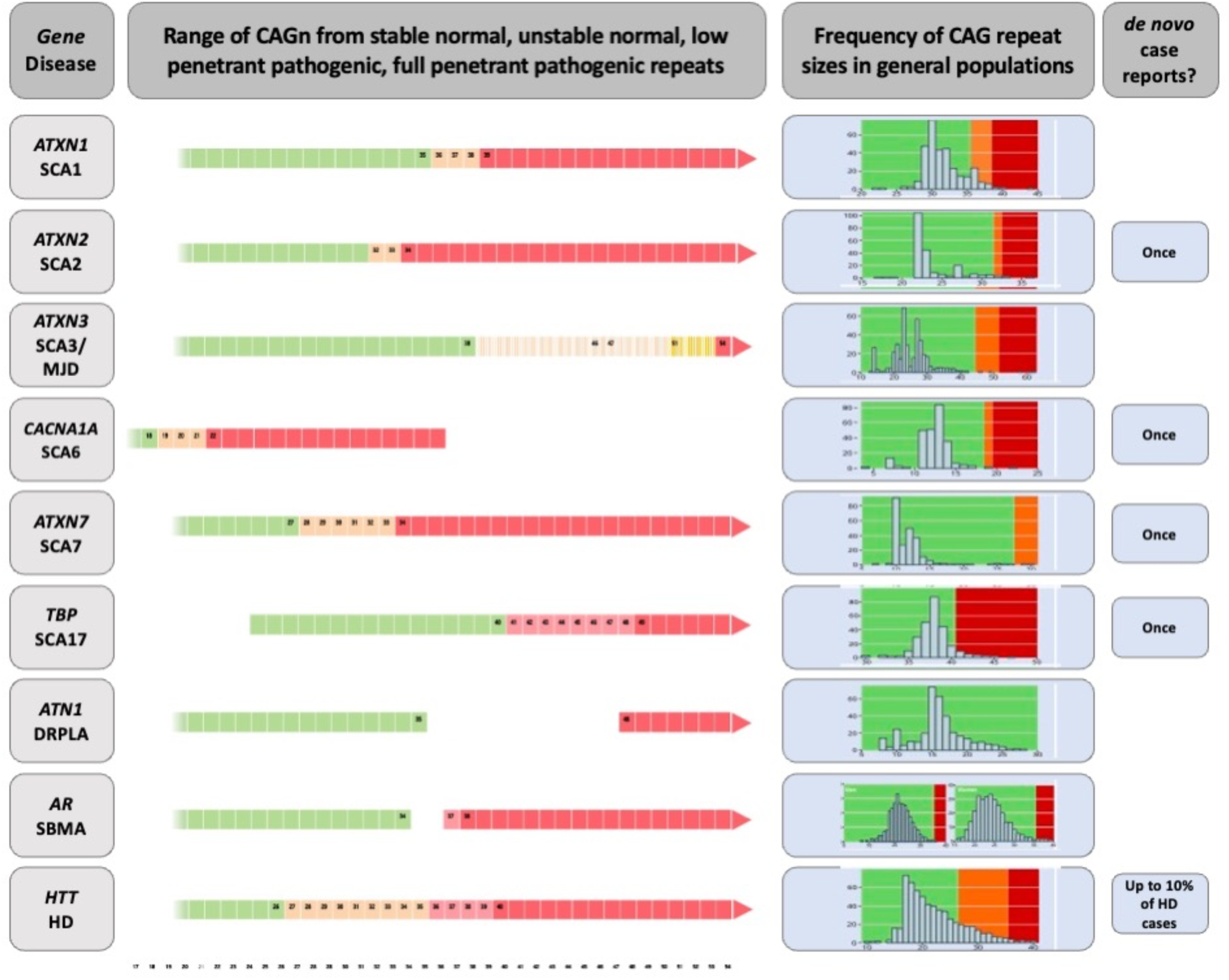
Diagram on the genetic characteristics of diseases related to polyglutamine expansions. The gene/disease column summarizes the disease-causing genes and the abbreviations of disease names of spinocerebellar ataxia types 1, 2, 3, 6, 7 and 17 (SCA1, SCA2, SCA3, SCA6, SCA7 and SCA17, dentatorubral-pallidoluysian atrophy (DRPLA), spinal and bulbar muscular atrophy (SBMA), and Huntington’s disease (HD). Data on CAGn presented in the second column was retrieved from Opal et al 1998, de Castilhos et al 2014, Gu et al 2004, Casey and Gomez 2019, Shizuka et al 1998, La Spada et al 2020, Mittal et al 2005, Toyoshima et al 2019, Bech et al 2010, Carroll et al 2018, La Spada et al 2017 and Caron et al 2020. The column called "Frequency of CAG repeat lengths in populations" shows the histograms of the alleles found in a normal population (data adapted from Gardiner et al 2019). Green means the range of normal alleles; orange, the range of intermediate alleles; and red, the range of pathological alleles.

PolyQ diseases are rare, progressive, and fatal, and share several clinical and genetic characteristics. Most are autosomal dominant diseases; the exception is SBMA, an X-linked disorder restricted to males due to the limited expression of the androgen receptor in females. Specific critical thresholds separate normal repeats from pathogenic ones (**Figure 1**). The age at onset (AO) or age at which the first neurologic manifestation was noted by the subject or their relatives, is usually in adulthood. Larger CAGexp lengths determine earlier AO and faster rates of disease progression. The CAGexp is unstable and tends to increase after meiosis; anticipation is quite frequent and might cause onset of symptoms in childhood. Clinical presentations of homozygous patients are not very different from those of heterozygotes. Most of these characteristics strongly support the hypothesis that the toxic and fully dominant gain of function underlies the pathogenesis. Reductions of the reproductive period per generation are expected in diseases associated with anticipation, which might drive many polyQ lineages to extinction. This poses the question of how many generations each lineage created by a *de novo* CAGexp expansion would last.

*ATXN2* and *ATXN3* are the genes related to SCA2 and SCA3/MJD, respectively. Both disorders are characterized by gait ataxia, pyramidal signs, a dystonic and/or rigid extrapyramidal syndrome, sensory losses, amyotrophy, and progressive external ophthalmoplegia (Pulst 2019; Paulson & Shakkottai 2020) [13,14]. AO, anticipation, neurologic manifestations, and survival after onset are similar across both diseases, so that they can be only distinguished with confidence by molecular testing (Bird et al 2019; Diallo et al 2018) [15,16].

There are some notable differences between both diseases. SCA3/MJD shows a relevant gap between the length of normal and expanded alleles (**Figure 1**), lacks *de novo* expansions, and preferentially segregates the expanded allele on meiosis. Evidence in favor of a few and old ancestral haplotypes have been obtained in several SCA3/MJD populations by robust studies (for instance, see Li et al 2019) [17]. *ATXN2* alleles show no gap between normal and expanded, *de novo* expansions have been described, and preferentially segregates the normal allele on meiosis (Sena et al 2021a and 2021b) [18, 19]. Reconstruction of ancestral haplotypes using single nucleotide polymorphisms (SNP) was restricted until recently to two markers and to the finding of a unique C-C haplotype worldwide (Choudhry et al 2001; Ramos et al 2008; Sonakar et al 2021) [20, 21, 22]. We have now used five SNPs and found at least eleven ancestral SCA2 haplotypes, just in South American families (Sena et al, submitted) [23]. These differences indicate that the biological contexts of *ATXN2* and *ATXN3* are quite diverse.

The similarities between SCA2 and SCA3/MJD clinical characteristics suggests that social and psychological impacts over their carriers should also be similar. This allows the presumption that the differences between SCA2 and SCA3/MJD transmissions to the offspring should not be attributed to distinct psychological or social pictures, but to the biological context of the expansions in *ATXN2* and *ATXN3*. Due to that, SCA2 and SCA3/MJD are probably good prototypes to test the dynamics of the CAGexp in general and to answer the question: "for how many generations will the descendants of a *de novo* expansion last?" This was the aim of the present study. The specific aims were to adapt a classical equation on allele dynamics in population genetics to be used in the case of dominant alleles related to late onset neurodegenerative disorders; and then, to test if the results of the model match with the existing epidemiological evidence on ancestral lineages and anticipation, in SCA2 and SCA3/MJD.

## 2. Subjects and methods

### 2.1 Subjects

The following human populations were used in this work: EUROSTAT (European Statistical Office) 2019 data [24] were used to establish fertility rates of normal women stratified by life year. Measures of fitness, segregation distortion, CAGexp instability and anticipation related to SCA2 or SCA3/MJD subjects were obtained from two meta-analyses published elsewhere (Sena et al. 2021a and 2021b) [18, 19]. Individual participant data (IPD) were obtained from other two original publications to estimate the reduction in AO attributable to each additional CAG repeat in CAGexp in SCA3/MJD (1,112 individuals) (de Mattos et al. 2018) [25], and in SCA2 (93 individuals) (Pereira et al 2015) [26]. Number of childbirths before or after age of onset of first symptom (AOfs) of 203 SCA2 and 334 SCA3/MJD carriers were obtained from two other IPD sources (Souza et al 2016; Sena et al 2019) [27, 28]. Finally, we used the stability/instability of the unexpanded allele with intermediate *ATXN2* length, observed in the general population and described in 57 individuals (Almaguer-Mederos et al 2018) [29].

### 2.2 Methods

To estimate the fate of CAGexp transmissions after a *de novo* expansion, one hypothetical expanded allele was assigned to a hypothetical founder of a lineage, and Monte Carlo methods were used to assign a random genotype to each descendant in several generations, based on segregation of the parental alleles, in a way like the gene dropping methodology (MacCluer et al 1986) [30]. The frequencies of the expanded alleles across generations, and the probabilities of extinction of these expanded alleles, were then assessed in a hypothetical population. The model assumed the absence of *de novo* mutations, genetic drift, and gene flow. In contrast, the effect of three mechanisms that could change the frequency of the expanded alleles were included: differential fitness of carriers of the expanded allele compared to non-carriers; transmission probabilities according to distortion in the segregation of the expanded allele; and anticipation.

#### 2.2.1 Adaptation of classical method on natural selection effect

The classical equation from population genetics theory encompasses fitness and distortion in allelic segregation as two selective forces of interest:

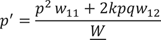

Where:

*p’* = Frequency of p allele in the subsequent generation
*p^2^* = Frequency of p allele in homozygotes
*w_11_* = Fitness of the p allele in homozygotes
*k* = segregation coefficient, where 0.5 represents Mendelian segregation.
*p* = frequency of the p allele
*q* = frequency of the q allele
*w_12_* = fitness of heterozygotes.
*W* = average fitness.

Adaptations were made to match the original equation to the specific characteristics of polyQs diseases, as follows.

First, an anticipation coefficient (*antcoeff)* was included to account for the influence of anticipation on the allele frequency at each generation of a lineage. The *antcoeff* ranged from zero to one, where zero corresponds to symptoms starting before the beginning of the fertile life (proposed as being 12 years of age) and one is related to symptoms that begin after the end of the fertile life (proposed as being 50 years of age). The extreme values represent the worst and the best scenarios for reproductive life, respectively, and the anticipation coefficient values were the mathematical expression of the relation between AOfs and the fertility rate of the ages’ interval arising from the new cutoff in the reproductive period imposed by the AOfs, in a given generation, in the studied lineage.

The next adjustments aimed to simplify the model. As the expanded alleles have a very low frequency, homozygosity is so rare that is practically non-existent: as *p²* ≈ 0, then it was removed from the equation. The expanded alleles have complete or near complete dominance, without any clear dosage effect when present in double dose (Sanpei et al 1996; Saute & Jardim 2015) [31, 32]. Since penetrance is close to 100%, the expanded allele cannot “protect” itself from the action of natural selection when it is in heterozygosity. This implies that the frequency of the expanded allele in the subsequent generation (*p’*) is essentially modulated by the intrinsic selective forces associated with *p* only, and not with *q* (the frequency of non-expanded allele). Due to that and to the fact that *q* frequency is close to 1, *q* was also removed from the equation.

The last adjustment was to directly use the relative *w* fitness into the equation instead of using the carriers’ *W* fitness divided by the general *W* fitness of the population.

With this, we arrived at:

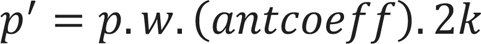

Where:

*p* = frequency of the expanded allele
*p’* = frequency of the expanded allele in the subsequent generation
*w* = relative fitness
*antcoeff* = anticipation coefficient
*k* = segregation coefficient

#### 2.2.2 The anticipation coefficient *antcoeff*

We arbitrated that the *antcoeff* would be equal to the average reproduction rate of a given generation, in the lineage of a common ancestor. Each generation of carriers would be prone to show modifications in their reproduction rate, due to the change of their average reproductive period, provoked, in turn, by the average anticipation of that generation.

We assume that each new anticipation will be associated with an additional reduction in the fertile period. Although symptomatic people might continue to have children during the early years of their illness, at some point, their clinical state will interfere with the reproductive capacity - either because of the children’s threats related to motor incapacity of a parent, or because of the reduced opportunities of sexual relationships required for reproduction. The conceptual relationship between anticipation and reproduction reduction has been studied and discussed elsewhere (Prestes et al 2008; Sena et al 2019) [28, 33]. In the datasets studied here, offspring of SCA2 and SCA3/MJD carriers were born at a mean (SD) of 13.69 (12.04) and 12.72 (12.84) years before the onset of symptoms, respectively; 92% and 91.7% of them were born before the onset of symptoms, in these diseases (data not shown). These birth rates that occurred after the onset of symptoms were then added to the value of the *antcoeff,* to bring them closer to reality of the reproduction rates.

To define how reductions in the reproductive period modify the anticipation coefficient in each generation, fertility rates in different age groups of a general population were established, using EUROSTAT data on the fertility of European women in 2019. In order to know how much each age group contributed to the overall fertility rate of that population in a given year, the total area below fertility function was calculated **(Figure S1 A)** and normalized to the value of 1, equal to the best anticipation coefficient, that was also equal to 1 when the disease onset is after the end of the reproductive period **(Figure S1 B)**. Each reduction in the reproductive period due to anticipation then reduced the area of the plot and reduced the value of the anticipation coefficient.

The next estimation to be made was that of the AOfs variation with each new generation, in order to extrapolate the corresponding reduction in the reproductive period.

All cohorts published to date were biased in favor of high anticipations (Sena et al 2021a and 2021b) [18, 19]. Therefore, the anticipation data retrieved from the literature was not directly used. Instead, we used a combination of two other pieces of information: the average instability of CAGexp transmission in each generation, and how much an increase of one CAG repeat reduces the AOfs. Studies on CAGexp transmissions were more representative since they commonly include asymptomatic as well as symptomatic individuals (Souza et al 2016; Sena et al 2019) [27, 28]. The product of [instability of transmission x AOfs-per-additional-CAGexp] estimated the anticipation in the subsequent generation. This multiplication produced smaller anticipations than the direct measurements; therefore, this procedure reduced distortions due to the literature bias in favor of excessive anticipations.

A linear regression performed for SCA3/MJD carriers from the IPD mentioned before (de Mattos et al 2018) [25] showed that each additional CAG repeat at expanded *ATXN3* was associated to a 1.652- year reduction in AOfs. To calculate the same for SCA2, we used the IPD from a publication on Brazilian, Peruvian, and Uruguayan individuals (Pereira et al 2015) [26]. Each additional CAGexp at *ATXN2* was related to a reduction of 1.877 years in AOfs (**Table 1**).

**Table 1.** Variables related to CAG repeats at *ATXN2* and *ATXN3*, used in the present adapted model. Data related to expanded repeats was obtained from heterozygous carriers; data related to normal repeats was obtained from non-carriers. Data is presented as means (standard deviation).

| Allele | Fitness,<br>w | Segregation<br>distortion,<br>k | Instability<br>during meiosis | AO reduction due to<br>each CAG repeat<br>added in the<br>expanded repeat |
| --- | --- | --- | --- | --- |
| ATXN2,<br>expanded<br>allele | 1.50<br>(0.25) *<br>Sena et al 2019 | 0.404<br>(0.085) *<br>Sena et al 2019 | 2.42<br>(5.655)<br>Sena et al 2021 | 1.877(1.86) |
| ATXN3,<br>expanded<br>allele | 1.45<br>(0.25) *<br>Prestes et al 2008 | 0.640<br>(0.085)<br>Sena et al 2021b | 1.23<br>(5.126)<br>Sena et al 2021b | 1.652(1.729) |
| ATXN2,<br>normal allele | 1.00 *<br>(0.25) * | 0.596<br>(0.085)<br>Sena et al 2019 | 0.23<br>(0.468)<br>Almaguer-Mederos et al<br>2018 |  |
| ATXN3,<br>normal allele | 1.00 *<br>(0.25) * | 0.360<br>(0.085)<br>Sena et al 2021b | 0.00 *<br>(0.468) * |  |

\* Imputed values. Reasons for each imputation were described in the text.

Contractions in the CAGexp repeat length can also occur, although very rarely documented (Cruz-Mariño et al 2014) [34]. After a contraction, the originally expanded allele of *ATXN2* and *ATXN3* might be transmitted as an unexpanded allele, causing the selective forces associated with SCA2 and SCA3/MJD diseases to no longer influence the dynamics of these descendant alleles. Although direct data were not available, contractions were taken into consideration in the resulting simulation model.

#### 2.2.3 Other variables to be included in the model

In addition to anticipation, the model needed to include fitness and segregation distortion, as mechanisms potentially associated with the long-term maintenance of polyQ diseases. Data on these forces were collected from previous systematic reviews (Sena et al 2021a and 2021b) [18, 19]. They are summarized in **Table 1**. Standard deviation (SD) values were needed to run the simulations. As there was no information about the SD of the segregation distortion in SCA2, the same SD found in SCA3/MJD segregation was assigned to the SCA2 model. Likewise, the SD of the fitness of both SCA2 and SCA3/MJD were lacking. In this case, the arbitrary value of 0.25 was imputed to them, as this value, although parsimonious, would allow for some overlap of individual fitness values between carriers and non-carriers.

Fitness, segregation rates and unstable transmissions associated with non-pathogenic alleles - those originated from contractions as well as the wildtype alleles - also need to be considered in the model. The fitness of the normal alleles is an a priori concept and is equal to one; the same SD of 0.25 were arbitrarily imputed to them. Transmission of intermediate-sized unexpanded *ATXN2* alleles was studied in the Cuban population (Almaguer-Mederos et al 2018) [29]. Studies on the transmission of the normal *ATXN3* alleles were not found; due to that, an arbitrary neutral value of zero with a standard deviation of 0.468, similar to found in the *ATXN2*, was assigned for the calculations. The data obtained from observational studies and used in the models were also systematized in **Table 1**.

### 2.3. Computer simulations on the dynamics of the expanded alleles

Means and SD of the variables considered so far, were used in the simulations on what occurs in the successive generations of a lineage. The emphasis on the use of SD was decided, in order that results could capture any potential scenario of real life. Thus, at each generational step, the simulated values brought the component of randomness to our results.

Measures of central tendency of descriptive variables were eventually presented as means (range) or as medians (range), according to the pattern of their distributions.

The simulations were performed in the R Statistical Package. The hypothetical initial frequency of the expanded allele was proposed to be 0.0001 for both *ATXN2* and *ATXN3* - sufficiently low to be considered plausible. The original ancestral expanded allele was proposed to correspond to the smallest length of the symptom-associated CAG repetitive sequences found in the datasets described in the section **2.1 Materials** - 34 and 54 CAGexp for SCA2 and SCA3/MJD, respectively (Sena et al 2019; de Mattos et al 2018) [25, 28]. Despite that, the model attributed pathogenicity to tracts with 34 CAG repeats in ATXN2 and with 51 CAG repeats in ATXN3, following the information described in **Figure 1**. The AOfs attributed for the first ancestor carrying this allele was the average AOfs found for the length of this expansion, in the same IPDs obtained from other studies - 55 and 65 years of age for SCA2 and SCA3/MJD, respectively (Sena et al 2019; de Mattos et al 2018) [25, 28].

At least 1,000 different lineages with allele frequencies randomly generated in each generation, considering the values described above, were simulated per CAGexp ancestor, covering a maximum of 650 generations. This number of generations was chosen because it corresponds to circa 16,250 years, or the approximate age of the oldest SCA3/MJD haplotype known so far, the TTACAC or Joseph lineage (Li et al 2019) [17]. Each of the 1,000 random lineages had their mean (SD) variables described above individually simulated in each generation. The values of the frequencies of the descendent alleles generated were re-entered into the equation to calculate the allele frequency in the subsequent generation. This process was carried out in loops until the frequency of the expanded allele reached 0 or 1 (fixation), in the possible descendent lines, or reached 650 generations - the proposed maximum observation time. The R source code to perform these simulations is described in the **Appendix.**

## 3. Results

### 3.1 Frequencies of the expanded allele at *ATXN2* across generations

From the 1,000 lineages simulated as descendants of the ancestral expansion of 34 repeats in *ATXN2*, 933 were eliminated in a median value of 10 generations, the extinction ranging between the 2nd and 121st generations. The frequency of alleles eliminated by generation and the number of generations that the allele remained in the population are represented in **Figure S2 A,** and **Figure 2A**.

**Figure 2.**
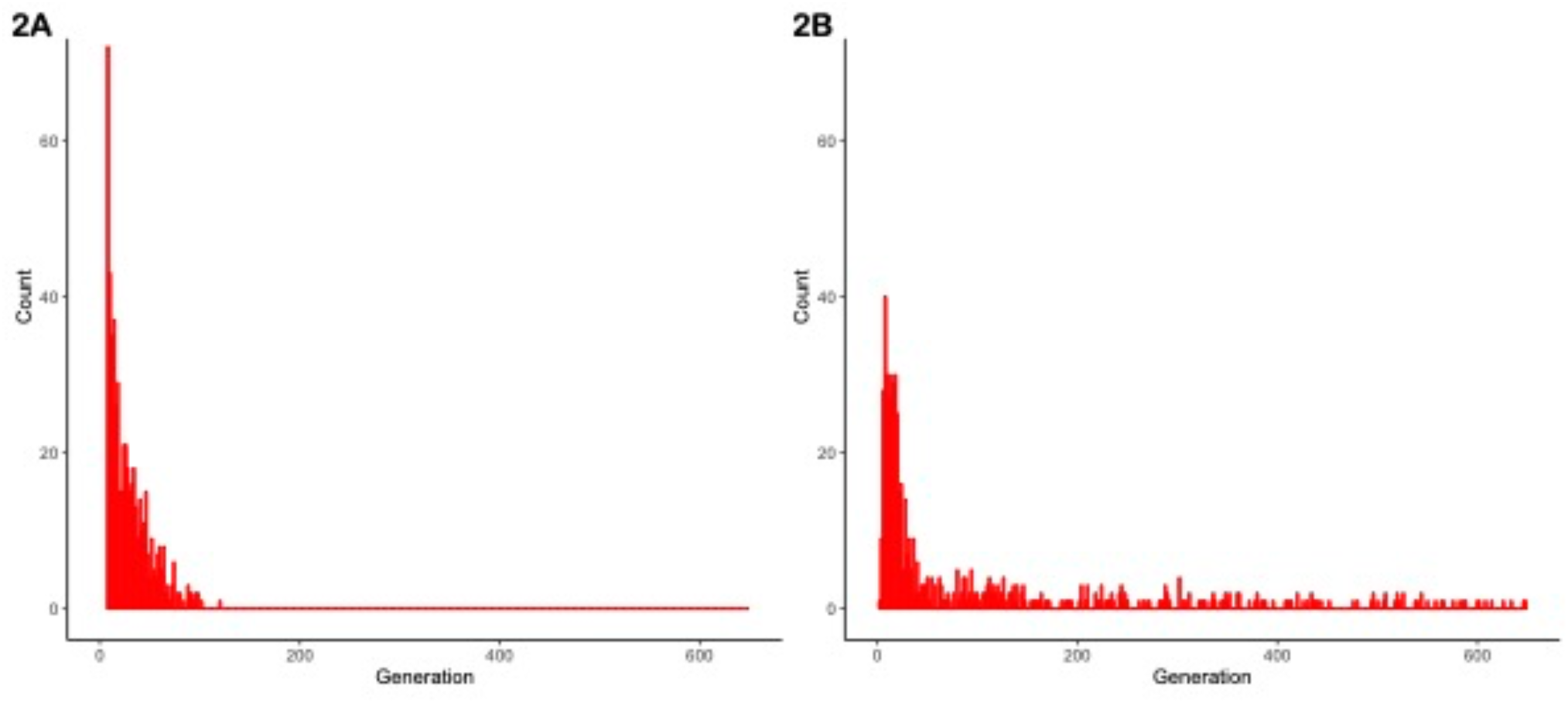
Fate of 1,000 lineages simulated as descendants of one ancestral with a CAG expansion and with an initial population frequency of 0.0001. (A) Proportion of descents with expanded repeats, per generation, after the first ancestor with 34 repeats in *ATXN2*. (B) Proportion of descents with expanded repeats, per generation, after the first ancestor with 54 repeats in *ATXN3*.

From these 1,000 simulated lineages, 67 were fixed after a median (range) of 60 (34 to 113) generations. In the generation in which the allele was fixed, the median (range) repeat length was of 32.00 (22.72 to 39.14) repeats - i.e., either non-pathogenic or borderline allele, in relation to SCA2 symptoms. The frequency of fixed alleles across generations are shown in **Figure 3A**. **Figure 3B** shows that all those 67 lineages in which the allele were fixed in the background population, turned out to be expanded, resulting in AOfs before the start of the reproductive period of life (**Figure 3C)**. These lineages would then become extinct in up to the 170th generation (**Table 2**).

**Figure 3.**
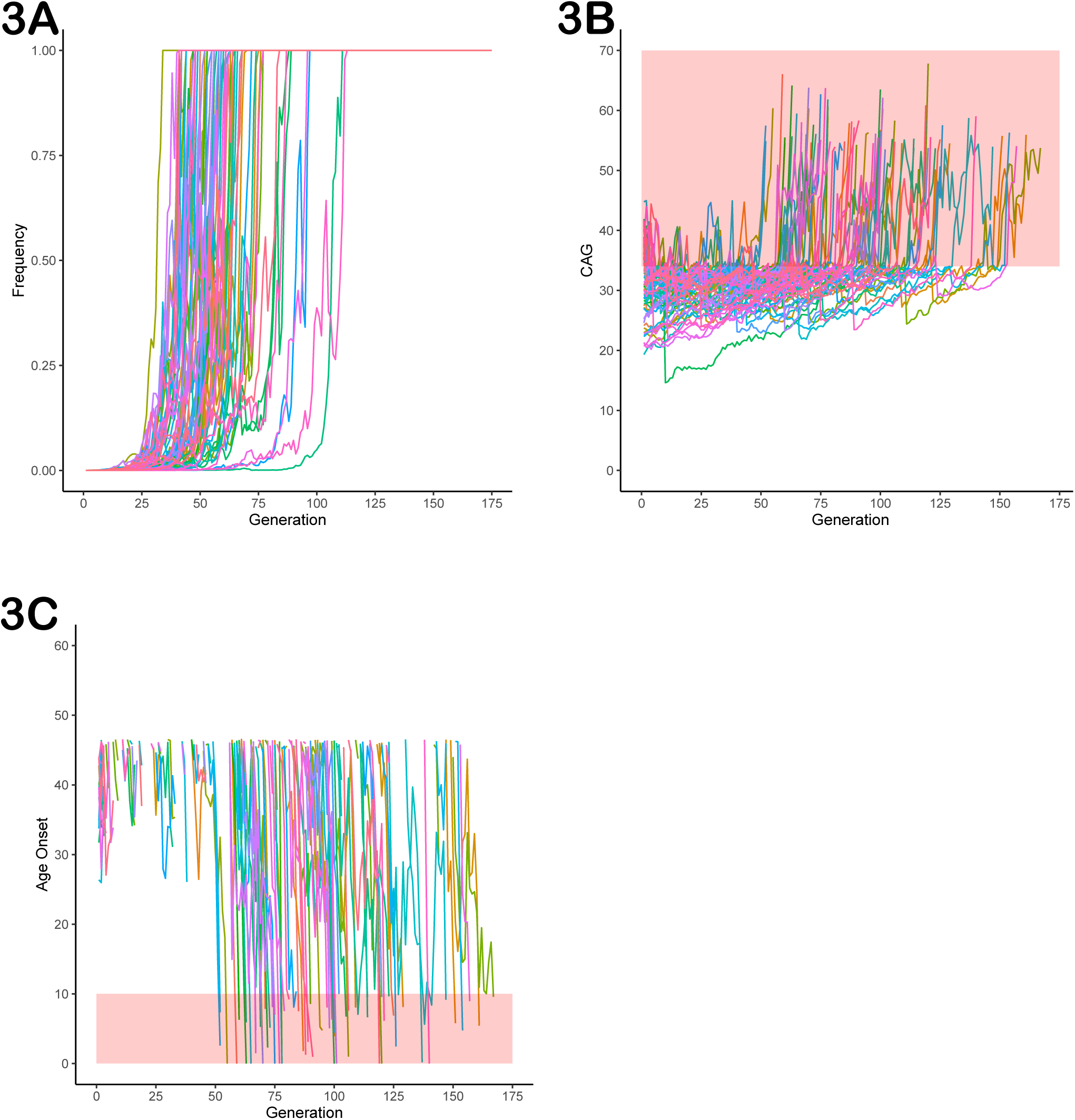
Data on the 67 lineages where fixed CAG expanded alleles at *ATXN2* were obtained after 1,000 simulations (simulations stopped when lineages were fixed). (A) Frequency of descendant alleles across generations until they were fixed. (B) The repeat lengths in the lineages where the descendant allele was fixed, per generation. The bold area in the graphic represents the pathological range of CAG repeat lenghts (or CAGexp). (C) The predicted mean age at onset of gait ataxia per generation of each fixed lineage. The bold area in the graphic represents ages of onset younger than 10 years old. The lines describing the AO in each lineage were interrupted at 46.55 years of age, at the top of the chart, since this was the AO predicted to be related to the 34 CAG repeats, the shortest expansion in the pathological range.

**Table 2.** Comparisons between the fates of *ATXN2* and *ATXN3* lineages produced by computer simulations from their expanded ancestors, in the 650th generation.

|  | Lineages<br>eliminated from<br>the population | Fixed alleles |  | Lineages that<br>remained in the<br>population |
| --- | --- | --- | --- | --- |
|  |  | Lineages extinct<br>after fixation | Lineages held as<br>fixed |  |
| *ATXN2* | 933 | 67 | 0 | 0 |
| *ATXN3* | 593 | 43 | 7 | 357 |
| *p* | p<0,001 | ( n.s) | (n.s) | p<0,001 |

### 3.2 Frequencies of the expanded allele at *ATXN3* across generations

From the 1,000 lineages simulated as descendants of the ancestral expansion with 54 repeats in *ATXN3*, 593 were eliminated in a median of 29 generations, the extinction ranging between 3 and 649 generations. The median size of the CAG repeats when the lineages were eliminated was 84, ranging from 39.45 to 88. The frequency of alleles eliminated by generation and the histogram of the number of generations where the allele was deleted are shown in **Figure 2B** and in **Figure S2 B.**

Of the same 1,000 simulated lineages, 50 were fixed and their frequencies across the generation are shown in **Figure 4A**. Fixation occurred at a median (range) of 19.50 (14 to 63) generations. **Figure 4B** shows the repeat lengths until the allele was fixed. In all the fixed lineages, the allele was expanded when it became fixed: they had a mean (range) of 64.64 (53.20 to 75.86) CAG repeats; the mean (range) AO of their carriers was 50.34 (14.98 to 70.41) years. After turning fixed, further instabilities continued to occur in the descendants (**Figure 4B).** Of the 50 fixed *ATXN3* lineages, the 43 that continued to expand were eliminated due to severe anticipations in AO (**Figure 4C**); the seven lineages transmitted after the 650th generation presented a contraction, carrying a limitrophe allele between normal and pathogenic CAG repeat lengths (**Figure 4B**).

**Figure 4.**
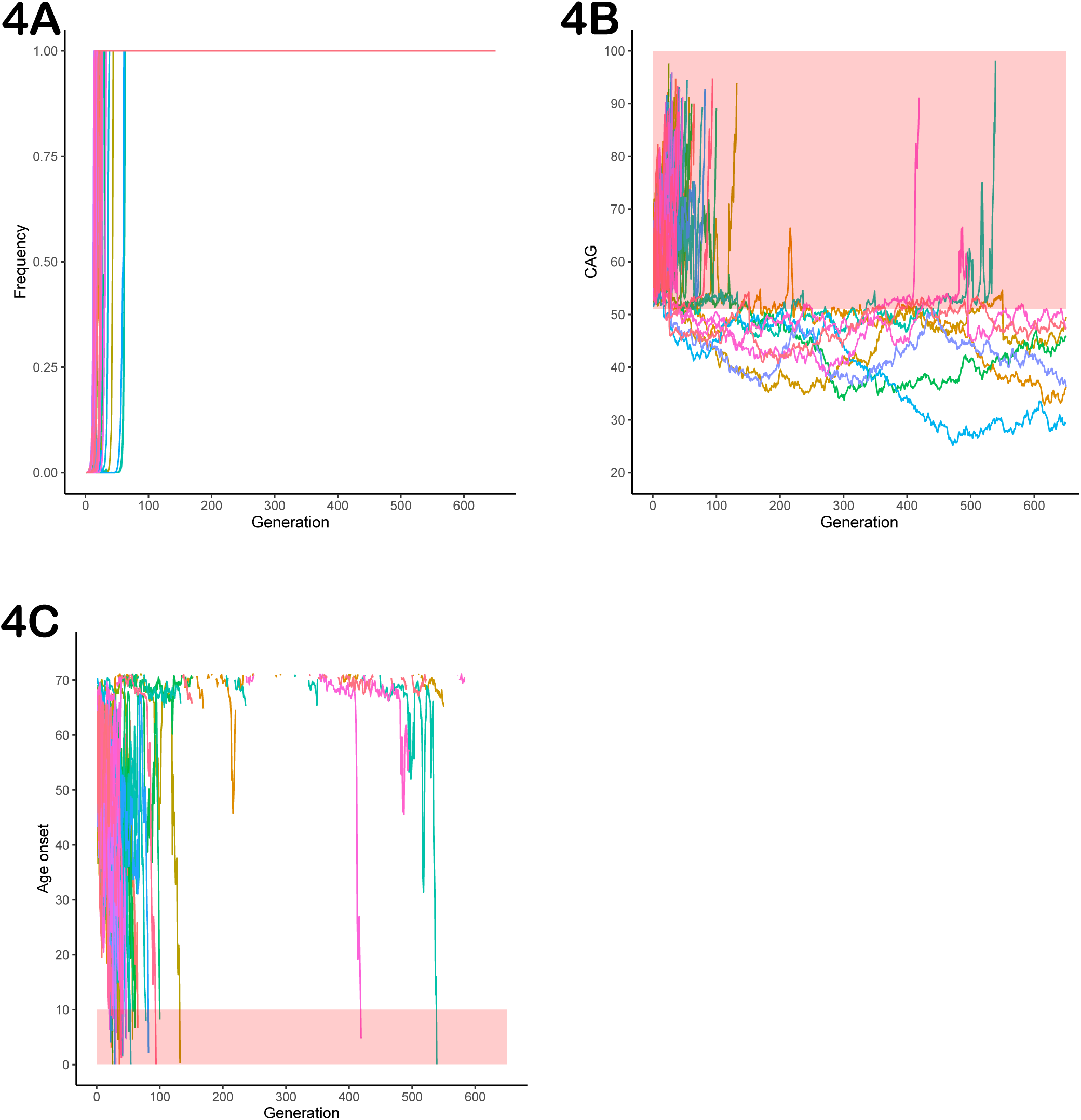
Data on the fifty lineages where fixed CAG expanded alleles at *ATXN3* were obtained after 1,000 simulations (simulations stopped when lineages were fixed). (A) Frequency of descendant alleles across generations until they were fixed. (B) The repeat lengths in the lineages where the descendant allele was fixed, per generation. The bold area in the graphic represents the pathological range of CAG repeat lengths (CAGexp). (C) The predicted mean age at onset of gait ataxia per generation of each fixed lineage. The bold area in the graphic represents ages of onset younger than 10 years old. The lines describing the AO in each lineage were interrupted at 71.14 years of age, at the top of the chart. They were related to CAG repeat sizes that were outside the pathological range, according to the regression.

Finally and more importantly, the 357 lineages where the simulated alleles were not eliminated nor fixed, had their frequencies across generations shown in **Figure 5A** and **Figure 5B**. These 357 lineages that remained in the population without fixed alleles showed CAG repeats transitioning between the expanded (equal or larger than 51 repeats) and non-expanded ranges (**Figure 5C**), suggesting that this phenomenon might make a lineage to survive for many centuries. Of note, the allele frequencies of the expanded repeats (51 repeats or more) remained very low and reached a median (IQR) 2.7e-107 (1.960994e-108) of in the 650th generation. When the 13 lineages that presented expanded alleles in generation 650 are presented separately, this alternation between the expanded and non-expanded allele can be more clearly observed (**Figure 5D**). **Table 2** summarizes these different fates of SCA3/MJD lineages and compares them to those of SCA2 lineages.

**Figure 5.**
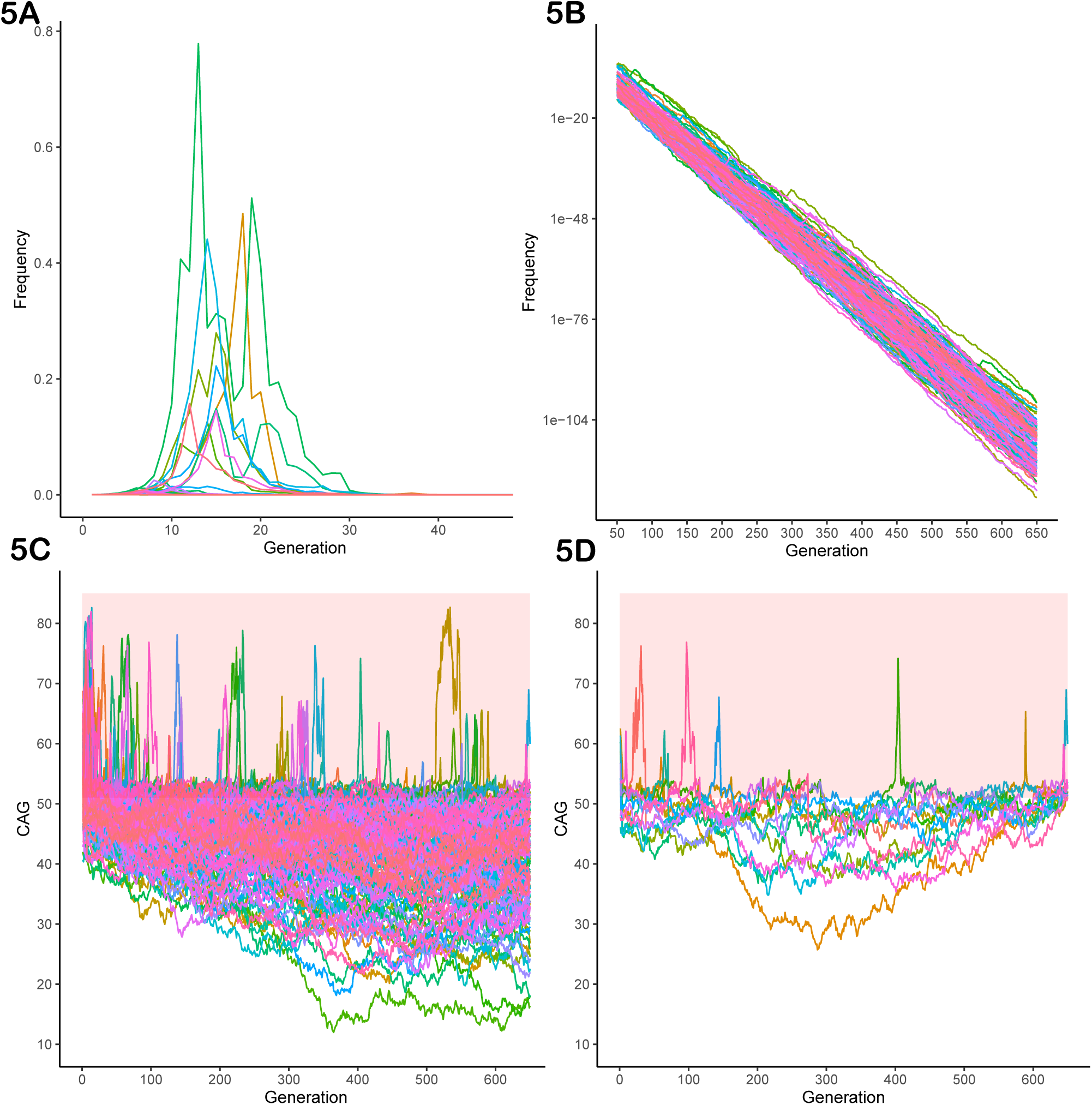
The 351 lineages of the CAG expanded alleles at *ATXN3* that remained non-fixed in the 1,000 simulations. (A) Frequency of descendant alleles until the 650th generation. Most have residual frequencies, very close to zero, and are not discernible on the graph. (B) The same in logarithmic scale. (C) The repeat lengths in these lineages, per generation. Note the transitions between the pathogenic and non-pathogenic CAG repeat lengths. The bold area in the graphic represents the pathological range of CAG repeat lengths (CAGexp). (D) The repeat lengths in the 13 lineages that presented expanded alleles in generation 650. The bold area in the graphic represents the pathological range of CAG repeat lengths (CAGexp). The repeat lengths alternate between the expanded and non-expanded ranges in generations that antecede the 650th one.

## 4. Discussion

Data on prevalence and anticipation have been difficult to put together in a unified biological explanation for polyQ diseases, as they are in contradiction with each other. Phenomena such as increased fitness and preferential segregation of the mutant alleles were then proposed to balance anticipation. But the empirical results, either because they were sparse or heterogeneous, kept the explanatory hypotheses in abeyance. The present model on the dynamics of expanded CAG alleles obtained compatible scenarios with current epidemiology, precisely using the empirical data on anticipation available at the moment. Our model suggests that these data are reliable and even sufficient to draft an acceptable and understandable explanation on why polyQ diseases might be present in populations for long periods of time.

The term "dynamics of the CAGexp" means the intrinsic, mutation-driven pattern of change in time of CAGexp which should be, by its nature, a multifactorial event. These dynamics might depend on the repeat motif itself (length, interruptions, etc), on the surrounding sequence, and on other factors that interplay with this surrounding context (sex and parental age, for instance) (Andrés et al 2022) [35]. To model the dynamics of the CAGexp, we have used computer simulation, adding to so many other applications related to population structure and evolutionary genetics (Hoban, Bertorelle & Gaggiotti 2012) [36].

Available empirical data on the evolutionary mechanisms at work on polyQs are certainly incomplete. Despite this, with data already available, the model predicted intergenerational dynamics that ended up being compatible with both apparently incoherent empirical data from SCA3/MJD - the few ancestral lineages with a long survival - and also the relatively more coherent facts associated with SCA2 - the serious anticipations due to dramatic expansions described in the literature and the multiple ancestral lineages (Sena et al, submitted) [23], both compatible with short survivals of its lineages.

In common, the CAGexp dynamics first went through an increase followed by a decrease in the frequencies of expanded alleles in successive generations. But the CAGexp alleles at *ATXN2* and *ATXN3* followed quite different trajectories (**Table 2**). The *ATXN3* allele might remain longer in the population, a fate due to the bias in favor of the expanded allele in the segregation of gametes, to favorable fitness, and to the less intense instability and anticipation than in relation to the expanded allele in *ATXN2* **(Figure S3).**

In contrast, the rise and fall of frequencies were quite sharp in SCA2. The model predicted that any real expansion in *ATXN2* would have a chance close to 98.6% of becoming extinct approximately 10 generations after its appearance. In this scenario, SCA2 recurrence in human populations distant as those of India, Cuba, and others, would depend upon *de novo* expansions.

The normal (CAG)_22_ allele in *ATXN2* is the most prevalent in the population (Laffita-Mesa et al 2012; Gardiner et al 2019) [37, 38]. CAG repeats at *ATXN2* are those with the lowest variance and allelic heterozygosity, between the *loci* related to polyQ diseases (Andrés et al 2002) [35]. These characteristics largely stem from the (CAG)_22_ allele being favored in meiotic segregation (Yu et al 2005, Chen et al 2013) [39, 40]. The internal sequence of this allele (CAG)_22_ contains two CAA interruptions, an important factor of stability in CAG repeats (Choudhry et al 2001) [20].

Given the positive selection of the (CAG)_22_ allele and the tendency of the expanded alleles to be rapidly withdrawn after they appear, SCA2 lineages should quickly disappear, and *de novo* expansions should be the most likely reason for the maintenance of SCA2 in populations. Our finding of at least eleven different ancestral SCA2 haplotypes among South American families are in line with this interpretation (Sena et al, submitted) [23]. This might be a case of mutation–selection balance. Intermediate or pre-expanded alleles associated with an unstable haplotype are the main sources of *de novo* expansions in HD (Warby et al 2009) [41]. The same might happen in SCA2. Indeed, an intermediate 32-repeat allele was detected in the asymptomatic father of a sporadic ataxic subject carrying 35 CAGexp in *ATXN2* (Futamura et al 1998) [42]. Although this was a unique report in the literature, it is worth reminding that *de novo* expansions most probably give rise to mild expansions and therefore to clinical manifestations very late in life, when parents are more frequently deceased and there is no way to document the phenomenon. Although the event might be very rare, it is necessary to clarify whether *de novo* expansions in *ATXN2* would come from predisposing haplotypes, with normal CAG repeats prone to instabilities and expansions when crossing meiosis. Comparisons among species suggest that C^rs695871^-C^rs695872^ (CC) is the oldest ancestral haplotype (Choudhry et al 2001) [20]; the (CAG)_22_ allele would have been raised more recently in humans, and would be associated with G^rs695871^-C^rs695872^ (GC) haplotype. Therefore, we can speculate that the *ATXN2* ancestral haplotype composed by GC haplotypes - and mostly with (CAG)_22_ - is associated with more stable repeats than those associated with the CC haplotype but analyzes with more markers are necessary to answer this question.

There is a continuum in the distribution of CAG alleles in *ATXN2* found in controls and in SCA2 carriers. The lack of a gap between normal and SCA2-associated alleles and the occurrence of *de novo* cases are peculiar characteristics that occur not only in SCA2 but also in other polyQ diseases, such as SCA6, SCA7, and HD (**Figure 1**). It is possible that the dynamics of the expanded allele of these other polyQs might be similar to that described here for SCA2.

In contrast, and as said before, SCA3/MJD is a polyQ disorder somehow different from most others, combining few ancestral haplotypes with a long-term permanence across generations (Martins et al 2007) [43]. These facts were in apparent contradiction with the serious anticipations registered in several SCA3/MJD cohorts, in such a way that one data or the other could be viewed with some doubt (Maciel et al 1995; Souza et al 2016) [22, 44]. Given its segregation distortion, fitness and *antcoeff* data, our mathematical model predicted that any expansion in *ATXN3* would have at least a 36.4% chance of lasting up to 650 generations. There was a substantial variability in the length of the transmitted repeats modeled here, and 13 of the 1,000 simulations reached the 650th generation as expanded. In fact, the mathematical model generated *ATXN3* lineages that appear to be able to stay indefinitely. We interpret that the segregation that favors the expanded *ATXN3* allele and the high fitness of SCA3/MJD are sufficient factors to explain the preservation of ancient ancestral haplotypes, such as the (at least) 16,000 years-old TTACAC or Joseph lineage (Martins and Sequeiros, 2018; Li et al 2019) [17, 45], until today. It is necessary to reflect that, although the actual ancient TTACAC alleles have not yet disappeared, it might be a matter of time for that to happen. In any case, our mathematical model not only supports the verisimilitude of the empirical data collected so far on SCA3/MJD, but also unifies them into a coherent set.

Our simulations have also revived an old hypothesis to explain the antiquity of the ancestral generations of SCA3/MJD: the existence of a haplotype that predisposes to expansions (Maciel et al. 1999) [46]. Although the TTACAC alleles carried by patients nowadays appear to have been passed on from a common ancestor for 650 generations - 16,000 years or more - this is not to say that ancestors with ataxic symptoms were frequent. **Figures 4C**, **5C** and **5D** denote that most of the expanded *ATXN3* lineages in the 650th generation previously oscillated between 37 and 50 CAG repeats, a range in which repeats do not produce symptoms. Although our model well supports the hypothesis that a haplotype might predispose for expansions, clinical and laboratory data published so far are less favorable, since SCA3/MJD carriers without a biological parent carrying one expansion have never been described in the literature. As far as we are aware, alleles in the range between normal and pathogenic have been described twice, but the authors did not clarify whether these alleles were non-penetrant (intermediate) or penetrant (pathogenic). Two subjects were detected in a collateral branch of a SCA3/MJD family, asymptomatic at 35 and 67 years, and carrying one *ATXN3* allele of 51 repeats (Maciel et al 2001) [47]. Another study measured the *ATXN3* CAG repeats in 16,547 subjects from five European population-based cohorts, and detected alleles with 46 and 49 CAG repeats (Gardiner et al 2019) [38]; the authors mentioned that they "had no long-term follow-up data on the participants", being "unable to confirm if carriers would have developed disease symptoms".

The lineages modeled here refer to the descendants of a first expansion carrier. It is interesting to consider the population environment as well, to understand the uncommon occurrence or even the lack of intermediate alleles in *ATXN3*. We have seen that short *ATXN3* alleles are transmitted preferentially in meiosis in the general population (Sena et al 2021a) [19]. This is the opposite of what happens in affected individuals, where the expanded allele is preferentially transmitted. Therefore, a disruptive selection takes place in *ATXN3*, that is, a selection in favor of extreme characteristics - favoring short alleles and expanded alleles and creating a gap between them. This gap would also explain the absence or ultra-rarity of intermediate alleles. In addition, it would also partially explain why *de novo* mutations have not been described in SCA3/MJD to date.

In any case, the fact that CAG tracts in *ATXN2*, *ATXN7* and *HTT*, among others, are prone to *de novo* expansion, while CAG tracts in *ATXN3* do not appear to be, needs to be further elucidated by observational and/or experimental studies. Discovering the reason for this discrepancy can have an impact even for future therapeutic or preventive management. One might suspect, for instance, that the CAG repeat of *ATXN3* has structural features in wild type alleles that confers a strong protection against instabilities. Or that the preferential segregation of the shortest allele in the presence of two normal alleles is a force to prevent a novel expansion of a normal allele originated from a contraction of an originally expanded allele.

To date, there is no indication whether the dynamics of the expanded allele in *ATXN3* finds similes among other polyQs. The best candidates to share these dynamics would be the polyQs for which no intermediate alleles were found, such as in DRPLA and in SBMA (**Figure 1**). In the case of SBMA, X-linked inheritance adds complexity to the description of intermediate alleles. DRPLA has additional similarities with SCA3/MJD: the gap between the normal range and the pathogenic range of CAGn is large (**Figure 1**); CAGexp on *ATN1* is favorably segregated (Ikeuchi et al 1996) [48]; there appear to be very few ancestral haplotypes (Martins et al 2003) [49]; and the existence of neurological subtypes, where manifestations are qualitative different, depending on AO.

Fixation of some descendant CAGexp alleles was a counterintuitive result of our model, but occurred in a minority of simulations, i.e., in 6.7% and 5% of *ATXN2* and *ATXN3*, respectively. The average CAG lengths of fixed lineages were 32 and 64 CAGexp for *ATXN2* and *ATXN3*, which are related to the late onset (or non-penetrant range, in *ATXN2* case) of the disease; part of them did not expand during fixation, thus not reducing the reproductive period of their carriers. When the simulations were continued, all fixed *ATXN2* lineages were eliminated due to the severe anticipation (**Figure 3C).** The 14 fixed *ATXN3* lineages at generation 650 had an unexpanded CAG repeat size; in some situations, their descents transited to the expanded range later **(Figure 4B)**. Indeed, Indeed, these scenarios seem highly hypothetical.

One can also speculate whether, in the past, other CAG repeats *loci* would have undergone pathogenic expansions causing maladaptive phenotypes. And that these alleles could have become extinct, so that the phenomenon would go unrecorded and be lost in human history.

Finally, as a theoretical work, the present study raised not direct evidence, but probabilities from existing empirical data on selective forces that converged with current epidemiology. It is worth emphasizing that we were interested in clarifying the effects of past history on present and not on future prevalences. Even so, further studies on prevalence and on ancestral haplotypes are still required to amplify these generalizations. Prevalence of SCA3/MJD in the Azores archipelago increased between 1981 and 2015 (de Araújo et al 2016) [50]. Similarly, prevalence of SCA1 in the Sakha (Yakut) people of Eastern Siberia increased between 1994 and 2013 (Platonov et al 2016) [51]. Monitoring the frequencies of polyQ diseases is relevant to keep track of eventual changes and to clarify the effectiveness of our model, in perspective.

In conclusion, the general dynamics of the CAGexp alleles seems to follow an increase in frequency for a few generations, followed by a decrease in frequency. Expanded *ATXN2* alleles showed a clear and rapid tendency to be eliminated from the population. Their maintenance in human populations must be explained by *de novo* expansions. To the contrary, expanded *ATXN3* alleles showed a tendency to remain longer in the population, a phenomenon explained at least by the favorable fitness, by the distortion in favor of the expanded allele or by a less intense instability and anticipation when compared to the expanded allele in *ATXN2*. These results contribute to the understanding of the survival of ancient origins for the *ATXN3* expansions. Finally, we think that the present mathematical model, combined with evidence of specific selective forces, can be used to simulate the dynamics of expanded alleles in other polyQ diseases.

## 5 Appendix

### Commands in R

A-To calculate the area that each age contributes to the fertility area trapz(area$age, area$"fertility_rate")

B-Transform the total area to a value of 1 and measure how much the reduction of 1 year of reproductive life reduces the total fertility rate

~~~
*area$area_s = 0*
*for(i in 2: dim(area)[1]){*
*age_s = area$age[i]*
*aux = subset(area, age <= age_s)*
*area$area_s[i] = trapz(aux$age, aux$"2019.0")}*
*area$areapad = area$area_s / trapz(area$age, area$"2019.0")*
~~~

3- To perform frequency simulations in subsequent generations

~~~
p <- NULL
cag_rep <- NULL
for(j in 1:n_linhagens){
   p1 <- *initial frequency*
   p2 <- NA
   cag_temp <- NA
   cag0 <- minor pathological cag
   print(paste0("lineage = ", n_linhagens))
   for(i in 1:generations){
~~~

cag<- cag0+ if(cag <minor pathological cag) {rnorm(n = 1, ***mean instability of the unexpanded allele***, standard deviation of unexpanded allele instability)} else {rnorm(n = 1, expanded allele mean instability, ***standard deviation of expanded allele instability***)}

~~~
    if(cag < ***minor pathological cag***){
       w <- rnorm(n = 1, ***fitness not affected***, ***fitness standard deviation unaffected***)
       k <- rnorm(n = 1, non-expanded allele segregation distortion, standard deviation of unexpanded allele segregation)
    } else {
       w <- rnorm(n = 1, ***fitness of those affected***, fitness standard deviation unaffected)
       k <- rnorm(n = 1,expanded allele segregation distortion,
***standard deviation of expanded allele segregation***)}

    if(w <= 0) w <- 0

antcoeff <- coefant$coef[i]

   ao <- -1.652 * cag + 155.389
   if(ao > 50){ antcoeff <- 1 }
   if(ao < 10){ antcoeff <- 0 }
   if(ao > 10 & ao < 50){ antcoeff <- predict(model, list(ao = ao), type="response") }
   cag0<-cag
   if(is.na(cag_temp)){
     cag_temp <- cag
    } else {
     cag_temp <- c(cag_temp, cag)
    }

      ao <- ***decrease in AO by CAG***\* cag + **(beta)**

      cag0<-cag
      if(is.na(cag_temp)){
         cag_temp <- cag
       } else {
          cag_temp <- c(cag_temp, cag)
       }
       if(antcoeff>1)antcoeff<-1
       print(paste0("generation ", i, " - cag length = ", cag, " - ant. coeff = ", antcoeff))
      

       if(p1 != 0 & p1 != 1){
          if(is.na(p2)){
             p2 <- p1*w*antcoeff*2*k
          } else {
             p2 <- c(p2, p1*w*antcoeff*2*k)
          }
        } else {
          if(p1 == 0) p2 <- c(p2, 0)
          if(p1 == 1) p2 <- c(p2, 1)
        }
        if(p2[i] >= 1){ p2[i] <- 1 }
        if(p2[i] <= 0){ p2[i] <- 0 }
        p1 <- p2[i]

        w_tab[j, i] <- w
        k_tab[j, i] <- k
        ao_tab[j, i] <- ao
        antcoeff_tab[j, i] <- antcoeff
       }
       p[[j]] <- p2
       cag_rep[[j]] <- cag_temp
   }
~~~

## 7. Author contributions

L.S.S., R.L. and L.B.J. contributed to the conception and design of the study; L.S.S., G.V.F., M.L.S.P. and L.B.J. contributed to the acquisition and analysis of data; L.S.S., R.L., and L.B.J. contributed to drafting the text and preparing the figures. All authors reviewed the manuscript.

## Supporting information

Sena et al, submitted (Related Article)

## Acknowledgements

This study was supported by Fundo de Incentivo à Pesquisa do Hospital de Clínicas de Porto Alegre (FIPE-HCPA) (grant numbers 2019-0254 and 2019-0169). LS, MLSP and LBJ were supported by Conselho Nacional de Desenvolvimento Científico e Tecnológico (CNPq), Brazil.

## 11. Supplemental Information

Figure S1 - Fertility rates of women from the European general population in 2019 (data from European Statistical Office). (A) Fertility rate per age. (B) Cumulative fertility rate.

Figure S2 -  Fate of the 1,000 lineages simulated as descendants of one ancestral with a CAG expansion with an initial population frequency of 0.0001. (A) Frequency of the descendant alleles of the original CAG expansion in *ATXN2* founder, per generation (B) Frequency of the descendant alleles of the original CAG expansion in *ATXN3* founder, per generation

Figure S3 -  The median (IQR) number of generations in which expanded alleles remained present in *ATXN2* and *ATXN3*, in gene dropping simulations.

## References

1. Adegbuyiroa A, Sedighia F, Pilkington AW, Groovera S, Legleitera J. Proteins containing expanded polyglutamine tracts and neurodegenerative disease. Biochemistry. 2017 March 07; 56(9): 1199–1217. doi:10.1021/acs.biochem.6b00936.

2. Bunting EL, Hamilton J, Tabrizi SJ. Polyglutamine diseases. Curr Opin Neurobiol. 2021 Sep 3;72:39-47. doi: 10.1016/j.conb.2021.07.001. Online ahead of print.PMID: 3448

3. Orr HT, Chung M-y, Banfi S, Kwiatkowski TJ, Servadio A, Beaudet AL, McCall AE, Duvick LA, Ranum LPW, Zoghbi HY. Expansion of an unstable trinucleotide CAG repeat in spinocerebellar ataxia type 1. Nat Genet. 1993; 4:221–226. [PubMed: 8358429]

4. La Spada, A. R., Wilson, E. M., Lubahn, D. B., Harding, A. E., Fischbeck, K. H. Androgen receptor gene mutations in X-linked spinal and bulbar muscular atrophy. Nature 352: 77–79, 1991. [PubMed: 2062380]

5. The Huntington’s Disease Collaborative Research Group. A Novel Gene Containing a Trinucleotide Repeat That Is Expanded and Unstable on Huntington’s Disease Chromosomes. Cell. 1993; 72:971–983. [PubMed: 8458085]

6. Kawaguchi Y, Okamoto T, Taniwaki M, Aizawa M, Inoue M, Katayama S, Kawakami H, Nakamura S, Nishimura M, Akiguchi I, Kimura J, Narumiya S, Kakizuka A. CAG expansions in a novel gene for Machado-Joseph disease at chromosome 14Q32.1. Nat Genet. 1994; 8:221–228. [PubMed: 7874163]

7. Komure O, Sano A, Nishino N, Yamauchi N, Ueno S, Kondoh K, Sano N, Takahashi M, Murayama N, Kondo I, Nagafuchi S, Yamada M, Kanazawa I. DNA analysis in hereditary Dentatorubral-Pallidoluysian Atrophy – correlation between CAG repeat length and phenotypic variation and the molecular-basis of anticipation. Neurology. 1995; 45:143–149. [PubMed: 7824105]

8. David G, Abbas N, Stevanin G, Durr A, Yvert G, Cancel G, Weber C, Imbert G, Saudou F, Antoniou E, Drabkin H, Gemmill R, Giunti P, Benomar A, Wood N, Ruberg M, Agid Y, Mandel JL, Brice A. Cloning of the SCA7 gene reveals a highly unstable CAG repeat expansion. Nat Genet. 1997; 17:65–70. [PubMed: 9288099]

9. Zhuchenko O, Bailey J, Bonnen P, Ashizawa T, Stockton DW, Amos C, Dobyns WB, Subramony SH, Zoghbi HY, Lee CC. Autosomal dominant cerebellar ataxia (SCA6) associated with small polyglutamine expansions in the alpha(1A)-voltage-dependent calcium channel. Nat Genet. 1997; 15:62–69. [PubMed: 8988170]

10. Koide R, Kobayashi S, Shimohata T, Ikeuchi T, Maruyama M, Saito M, Yamada M, Takahashi H, Tsuji S. A neurological disease caused by an expanded CAG trinucleotide repeat in the TATA-binding protein gene: a new polyglutamine disease? Hum Mol Genet. 1999; 8:2047–2053. [PubMed: 10484774]

11. Pulst SM, Santos N, Wang D, Yang HY, Huynh D, Velazquez L, Figueroa KP. Spinocerebellar ataxia type 2: polyQ repeat variation in the CACNA1A calcium channel modifies age of onset. Brain. 2005; 128:2297–2303. [PubMed: 16000334]

12. Margolis RL, Rudnicki DD, Holmes SE. Huntington’s disease like-2: review and update. Acta Neurol Taiwan. 2005; 14:1–8. [PubMed: 15835282]

13. Pulst SM. Spinocerebellar Ataxia Type 2. 1998 Oct 23 [Updated 2019 Feb 14]. In: Adam MP, Ardinger HH, Pagon RA, et al., editors. GeneReviews® [Internet]. Seattle (WA): University of Washington, Seattle; 1993–2022. Available from: https://www.ncbi.nlm.nih.gov/books/NBK1275/

14. Paulson H, Shakkottai V. Spinocerebellar Ataxia Type 3. 1998 Oct 10 [Updated 2020 Jun 4]. In: Adam MP, Ardinger HH, Pagon RA, et al., editors. GeneReviews® [Internet]. Seattle (WA): University of Washington, Seattle; 1993–2022. Available from: https://www.ncbi.nlm.nih.gov/books/NBK119

15. Bird TD. Hereditary Ataxia Overview. 1998 Oct 28 [Updated 2019 Jul 25]. In: Adam MP, Ardinger HH, Pagon RA, et al., editors. GeneReviews® [Internet]. Seattle (WA): University of Washington, Seattle; 1993–2022. Available from: https://www.ncbi.nlm.nih.gov/books/NBK1138/

16. Diallo A, Jacobi H, Cook A, Labrum R, Durr A, Brice A, Charles P, Marelli C, Mariotti C, Nanetti L, Panzeri M, Rakowicz M, Sobanska A, Sulek A, Schmitz-Hübsch T, Schöls L, Hengel H, Melegh B, Filla A, Antenora A, Infante J, Berciano J, van de Warrenburg BP, Timmann D, Boesch S, Pandolfo M, Schulz JB, Bauer P, Giunti P, Kang JS, Klockgether T, Tezenas du Montcel S. Survival in patients with spinocerebellar ataxia types 1, 2, 3, and 6 (EUROSCA): a longitudinal cohort study. Lancet Neurol. 2018 Apr;17(4):327–334. doi: 10.1016/S1474-4422(18)30042-5.

17 Li T, Martins S, Peng Y, Wang P, Hou X, Chen Z, Wang C, Tang Z, Qiu R, Chen C, Hu Z, Xia K, Tang B, Sequeiros J, Jiang H.Is the High Frequency of Machado-Joseph Disease in China Due to New Mutational Origins? Front Genet. 2019 Feb 20;9:740.

18. Sena LS, Dos Santos Pinheiro J, Hasan A, Saraiva-Pereira ML, Jardim LB. Spinocerebellar ataxia type 2 from an evolutionary perspective: Systematic review and meta-analysis. Clin Genet. 2021a Sep;100(3):258–267. doi: 10.1111/cge.13978. Epub 2021 May 27. PMID: 33960424.

19. Sena LS, Dos Santos Pinheiro J, Saraiva-Pereira ML, Jardim LB. Selective forces acting on spinocerebellar ataxia type 3/Machado-Joseph disease recurrency: A systematic review and meta-analysis. Clin Genet. 2021b Mar;99(3):347–358. doi: 10.1111/cge.13888. Epub 2020 Dec 2. PMID: 33219521.

20 Choudhry S, Mukerji M, Srivastava AK, Jain S, Brahmachari SK. CAG repeat instability at SCA2 locus: anchoring CAA interruptions and linked single nucleotide polymorphisms. Hum Mol Genet. 2001;10: 2437–2446

21 Ramos EM, Martins S, Alonso I, Emmel VE, Saraiva-Pereira ML, Jardim LB, Coutinho P, Sequeiros J, Silveira I. Common origin of pure and interrupted repeat expansions in spinocerebellar ataxia type 2 (SCA2). Am J Med Genet B Neuropsychiatr Genet. 2010 Mar 5;153B(2):524–531. doi: 10.1002/ajmg.b.31013. PMID: 19676102.

22 Sonakar AK, Shamim U, Srivastava MP, Faruq M, Srivastava AK. (2021). SCA2 in the Indian population: Unified haplotype and variable phenotypic patterns in a large case series. Parkinsonism Relat Disord. doi: 10.1016/j.parkreldis.2021.07.011. Jul 14. PMID: 34298214.

23 Sena LS, Furtado GV, Fagundes NJR, Pedroso JL, Barsottini O, Ribeiro P, Cornejo-Olivas M, Vargas FR, Godeiro C, Medeiros PFV, Torales MBPP, Braga-Neto P, Camejo C, Jardim LB, Saraiva-Pereira, on behalf of Rede Neurogenetica. Spinocerebellar ataxia type 2 has multiple ancestral origins. Submitted.

24. https://ec.europa.eu/eurostat/databrowser/view/demo_find/default/table?lang=en

25. de Mattos EP, Kolbe Musskopf M, Bielefeldt Leotti V, Saraiva-Pereira ML, Jardim LB. Genetic risk factors for modulation of age at onset in Machado-Joseph disease/spinocerebellar ataxia type 3: a systematic review and meta-analysis. J Neurol Neurosurg Psychiatry. 2019 Feb;90(2):203–210. doi: 10.1136/jnnp-2018-319200. Epub 2018 Oct 18. PMID: 30337442.

26. Pereira FS, Monte TL, Locks-Coelho LD, Silva AS, Barsottini O, Pedroso JL, Cornejo-Olivas M, Mazzetti P, Godeiro C, Vargas FR, Lima MA, van der Linden H Jr, Toralles MB, Medeiros PF, Ribeiro E, Braga-Neto P, Salarini D, Castilhos RM, Saraiva-Pereira ML, Jardim LB; Rede Neurogenetica. ATXN3, ATXN7, CACNA1A, and RAI1 Genes and Mitochondrial Polymorphism A10398G Did Not Modify Age at Onset in Spinocerebellar Ataxia Type 2 Patients from South America. Cerebellum. 2015 Dec;14(6):728–30. doi: 10.1007/s12311-015-0666-8. PMID: 25869926.

27. Souza GN, Kersting N, Krum-Santos AC, Santos AS, Furtado GV, Pacheco D, Gonçalves TA, Saute JA, Schuler-Faccini L, Mattos EP, Saraiva-Pereira ML, Jardim LB. Spinocerebellar ataxia type 3/Machado-Joseph disease: segregation patterns and factors influencing instability of expanded CAG transmissions. Clin Genet. 2016 Aug;90(2):134–40. doi: 10.1111/cge.12719. Epub 2016 Feb 3. PMID: 26693702.

28. Sena LS, Castilhos RM, Mattos EP, Furtado GV, Pedroso JL, Barsottini O, de Amorim MMP, Godeiro C, Pereira MLS, Jardim LB. Selective Forces Related to Spinocerebellar Ataxia Type 2. Cerebellum. 2019 Apr;18(2):188–194. doi: 10.1007/s12311-018-0977-7. Erratum in: Cerebellum. 2018 Nov 19;:. Pereira MLS [corrected to Saraiva-Pereira ML]. PMID: 30219976.

29. Almaguer-Mederos LE, Mesa JML, González-Zaldívar Y, Almaguer-Gotay D, Cuello-Almarales D, Aguilera-Rodríguez R, Falcón NS, Gispert S, Auburger G, Velázquez-Pérez L. Factors associated with ATXN2 CAG/CAA repeat intergenerational instability in Spinocerebellar ataxia type 2. Clin Genet. 2018 Oct;94(3-4):346–350. doi: 10.1111/cge.13380. Epub 2018 Jun 29. PMID: 29756284.

30. MacCluer J.W., VandeBerg J.L., Read B. and Ryder O.A. (1986). Pedigree analysis by computer simulation. Zoo Biol. 5(2), 147–160.

31. Sanpei K, Takano H, Igarashi S, Sato T, Oyake M, Sasaki H, Wakisaka A, Tashiro K, Ishida Y, Ikeuchi T, Koide R, Saito M, Sato A, Tanaka T, Hanyu S, Takiyama Y, Nishizawa M, Shimizu N, Nomura Y, Segawa M, Iwabuchi K, Eguchi I, Tanaka H, Takahashi H, Tsuji S. Identification of the spinocerebellar ataxia type 2 gene using a direct identification of repeat expansion and cloning technique, DIRECT. Nat Genet. 1996;14:277–84

32. Saute JAM, Jardim LB. Machado Joseph disease: clinical and genetic aspects, and current treatment, Expert Opinion on Orphan Drugs, 2015; 3: 517–535, DOI: 10.1517/21678707.2015.1025747

33. Prestes PR, Saraiva-Pereira ML, Silveira I, Sequeiros J, Jardim LB. Machado-Joseph disease enhances genetic fitness: a comparison between affected and unaffected women and between MJD and the general population. Ann Hum Genet. 2008 Jan;72(Pt 1):57–64. doi: 10.1111/j.1469-1809.2007.00388.x. Epub 2007 Aug 7. PMID: 17683516.

34. Cruz-Mariño T, Laffita-Mesa JM, Gonzalez-Zaldivar Y, Velazquez-Santos M, Aguilera-Rodriguez R, Estupinan-Rodriguez A, Vazquez-Mojena Y, Macleod P, Paneque M, Velazquez-Perez L. Large normal and intermediate alleles in the context of SCA2 prenatal diagnosis. J Genet Couns. 2014 Feb;23(1):89–96. doi: 10.1007/s10897-013-9615-1. Epub 2013 Jun 28. PMID: 23813298.

35. Andrés AM, Lao O, Soldevila M, Calafell F, Bertranpetit J. Dynamics of CAG repeat loci revealed by the analysis of their variability. Hum Mutat. 2003 Jan;21(1):61–70. doi: 10.1002/humu.10151.

36. Hoban, S., Bertorelle, G. & Gaggiotti, O. Computer simulations: tools for population and evolutionary genetics. Nat Rev Genet 13, 110–122 (2012). 10.1038/nrg3130

37. Laffita-Mesa JM, Velázquez-Pérez LC, Santos Falcón N, et al. Unexpanded and intermediate CAG polymorphisms at the SCA2 locus (ATXN2) in the Cuban population: evidence about the origin of expanded SCA2 alleles. European Journal of Human Genetics. 2012;20(1):41–49. doi:10.1038/ejhg.2011.

38. Gardiner SL, Boogaard MW, Trompet S, et al. Prevalence of Carriers of Intermediate and Pathological Polyglutamine Disease-Associated Alleles Among Large Population-Based Cohorts. JAMA Neurol. 2019;76(6):650–656. doi:10.1001/jamaneurol.2019.0423

39. Yu F, Sabeti PC, Hardenbol P, et al. Positive selection of a pre-expansion CAG repeat of the human SCA2 gene. PLoS Genet. 2005;1(3):e41. doi:10.1371/journal.pgen.0010041

40. Chen XC, Sun H, Zhang CJ, Zhang Y, Lin KQ, Yu L, Shi L, Tao YF, Huang XQ, Chu JY, Yang ZQ. Positive selection of CAG repeats of the ATXN2 gene in Chinese ethnic groups. J Genet Genomics. 2013 Oct 20;40(10):543–8. doi: 10.1016/j.jgg.2013.08.003. Epub 2013 Sep 24. PMID: 24156920.

41. Warby SC, Montpetit A, Hayden AR, Carroll JB, Butland SL, Visscher H, Collins JA, Semaka A, Hudson TJ, Hayden MR. CAG expansion in the Huntington disease gene is associated with a specific and targetable predisposing haplogroup. Am J Hum Genet. 2009 Mar;84(3):351–66. doi: 10.1016/j.ajhg.2009.02.003. Epub 2009 Feb 26. PMID: 19249009; PMCID: PMC2668007.

42. Futamura N, Matsumura R, Fujimoto Y, Horikawa H, Suzumura A, Takayanagi T. CAG repeat expansions in patients with sporadic cerebellar ataxia. Acta Neurol Scand. 1998 Jul;98(1):55–9. doi: 10.1111/j.1600-0404.1998.tb07378.x. PMID: 9696528.

43. Martins S, Calafell F, Gaspar C, Wong VC, Silveira I, Nicholson GA, Brunt ER, Tranebjaerg L, Stevanin G, Hsieh M, Soong BW, Loureiro L, Dürr A, Tsuji S, Watanabe M, Jardim LB, Giunti P, Riess O, Ranum LP, Brice A, Rouleau GA, Coutinho P, Amorim A, Sequeiros J. Asian origin for the worldwide-spread mutational event in Machado-Joseph disease. Arch Neurol. 2007 Oct;64(10):1502–8.

44. Maciel P, Gaspar C, DeStefano AL, Silveira I, Coutinho P, Radvany J, Dawson DM, Sudarsky L, Guimarães J, Loureiro JE, et al. Correlation between CAG repeat length and clinical features in Machado-Joseph disease. Am J Hum Genet. 1995 Jul;57(1):54–61. PMID: 7611296; PMCID: PMC1801255.

45. Martins S, Sequeiros J. Origins and Spread of Machado-Joseph Disease Ancestral Mutations Events. Adv Exp Med Biol. 2018; 1049: 243–254.

46. Maciel P, Gaspar C, Guimarães L, Goto J, Lopes-Cendes I, Hayes S, Arvidsson K, Dias A, Sequeiros J, Sousa A, Rouleau GA.Study of three intragenic polymorphisms in the Machado-Joseph disease gene (MJD1) in relation to genetic instability of the (CAG)n tract. Eur J Hum Genet. 1999 Feb-Mar;7(2):147–56. doi: 10.1038/sj.ejhg.5200264.

47. Maciel P, Costa MC, Ferro A, Rousseau M, Santos CS, Gaspar C, Barros J, Rouleau GA, Coutinho P, Sequeiros J.Improvement in the molecular diagnosis of Machado-Joseph disease. Arch Neurol. 2001 Nov;58(11):1821–7. doi: 10.1001/archneur.58.11.1821.

48. Ikeuchi T, Igarashi S, Takiyama Y, et al. Non-Mendelian transmission in dentatorubral-pallidoluysian atrophy and Machado-Joseph disease: the mutant allele is preferentially transmitted in male meiosis. Am J Hum Genet. 1996;58(4):730–733.

49. Martins S, Matamá T, Guimarães L, Vale J, Guimarães J, Ramos L, Coutinho P, Sequeiros J, Silveira I. Portuguese families with dentatorubropallidoluysian atrophy (DRPLA) share a common haplotype of Asian origin. Eur J Hum Genet. 2003 Oct;11(10):808–11. doi: 10.1038/sj.ejhg.5201054. PMID: 14512972.

50. de Araújo MA, Raposo M, Kazachkova N, Vasconcelos J, Kay T, et al. (2016) Trends in the Epidemiology of Spinocerebellar Ataxia Type 3/Machado-Joseph Disease in the Azores Islands, Portugal. JSM Brain Sci 1(1): 1001.

51. Platonov FA, Tyryshkin K, Tikhonov DG, et al. Genetic fitness and selection intensity in a population affected with high-incidence spinocerebellar ataxia type 1. Neurogenetics. 2016;17(3):179–185. doi:10.1007/s10048-016-0481-5

52. Opal P, Ashizawa T. Spinocerebellar Ataxia Type 1. 1998 Oct 1 [Updated 2017 Jun 22]. In: Adam MP, Ardinger HH, Pagon RA, et al., editors. GeneReviews® [Internet]. Seattle (WA): University of Washington, Seattle; 1993–2022. Available from: https://www.ncbi.nlm.nih.gov/books/NBK1184/

53. de Castilhos RM, Furtado GV, Gheno TC, Schaeffer P, Russo A, Barsottini O, Pedroso JL, Salarini DZ, Vargas FR, de Lima MA, Godeiro C, Santana-da-Silva LC, Toralles MB, Santos S, van der Linden H Jr, Wanderley HY, de Medeiros PF, Pereira ET, Ribeiro E, Saraiva-Pereira ML, Jardim LB; Rede Neurogenetica. Spinocerebellar ataxias in Brazil--frequencies and modulating effects of related genes. Cerebellum. 2014 Feb;13(1):17–28. doi: 10.1007/s12311-013-0510-y. PMID: 23943520.

54. Gu W, Ma H, Wang K, et al. The shortest expanded allele of the MJD1 gene in a Chinese MJD kindred with autonomic dysfunction. Eur Neurol 2004; 52:107–11

55. Casey HL, Gomez CM. Spinocerebellar Ataxia Type 6. 1998 Oct 23 [Updated 2019 Nov 21]. In: Adam MP, Ardinger HH, Pagon RA, et al., editors. GeneReviews® [Internet]. Seattle (WA): University of Washington, Seattle; 1993–2022. Available from: https://www.ncbi.nlm.nih.gov/books/NBK1140/

56. Shizuka M, Watanabe M, Ikeda Y, Mizushima K, Okamoto K, Shoji M. Molecular analysis of a de novo mutation for spinocerebellar ataxia type 6 and (CAG)n repeat units in normal elder controls. J Neurol Sci. 1998 Nov 26;161(1):85–7. doi: 10.1016/s0022-510x(98)00270-6. PMID: 9879686.

57. La Spada AR. Spinocerebellar Ataxia Type 7. 1998 Aug 27 [Updated 2020 Jul 23]. In: Adam MP, Ardinger HH, Pagon RA, et al., editors. GeneReviews® [Internet]. Seattle (WA): University of Washington, Seattle; 1993–2022. Available from: https://www.ncbi.nlm.nih.gov/books/NBK1256/

58. Mittal U, Roy S, Jain S, Srivastava AK, Mukerji M. Post-zygotic de novo trinucleotide repeat expansion at spinocerebellar ataxia type 7 locus: evidence from an Indian family. J Hum Genet. 2005;50:155–7.

59. Toyoshima Y, Onodera O, Yamada M, et al. Spinocerebellar Ataxia Type 17. 2005 Mar 29 [Updated 2019 Sep 12]. In: Adam MP, Ardinger HH, Pagon RA, et al., editors. GeneReviews® [Internet]. Seattle (WA): University of Washington, Seattle; 1993–2022. Available from: https://www.ncbi.nlm.nih.gov/books/NBK1438/

60. Bech S, Petersen T, Nørremølle A, Gjedde A, Ehlers L, Eiberg H, Hjermind LE, Hasholt L, Lundorf E, Nielsen JE. Huntington’s disease-like and ataxia syndromes: identification of a family with a de novo SCA17/TBP mutation. Parkinsonism Relat Disord. 2010 Jan;16(1):12–5. doi: 10.1016/j.parkreldis.2009.06.006. PMID: 19595623.

61. Carroll LS, Massey TH, Wardle M, Peall KJ. Dentatorubral-pallidoluysian Atrophy: An Update. Tremor Other Hyperkinet Mov (N Y). 2018;8:577. Published 2018 Oct 1. doi:10.7916/D81N9HST

62. La Spada A. Spinal and Bulbar Muscular Atrophy. 1999 Feb 26 [Updated 2017 Jan 26]. In: Adam MP, Ardinger HH, Pagon RA, et al., editors. GeneReviews® [Internet]. Seattle (WA): University of Washington, Seattle; 1993–2022. Available from: https://www.ncbi.nlm.nih.gov/books/NBK1333/

63. Caron NS, Wright GEB, Hayden MR. Huntington Disease. 1998 Oct 23 [Updated 2020 Jun 11]. In: Adam MP, Ardinger HH, Pagon RA, et al., editors. GeneReviews® [Internet]. Seattle (WA): University of Washington, Seattle; 1993–2022. Available from: https://www.ncbi.nlm.nih.gov/books/NBK1305/

