## Supplementary material for "A model for the dynamics of expanded CAG repeat alleles: *ATXN2* and *ATXN3* as prototypes": Figure S1, S2 and S3

**Figure S1.** Fertility rates of women from the European general population in 2019 (data from European Statistical Office). (A) Fertility rate per age. (B) Cumulative fertility rate.

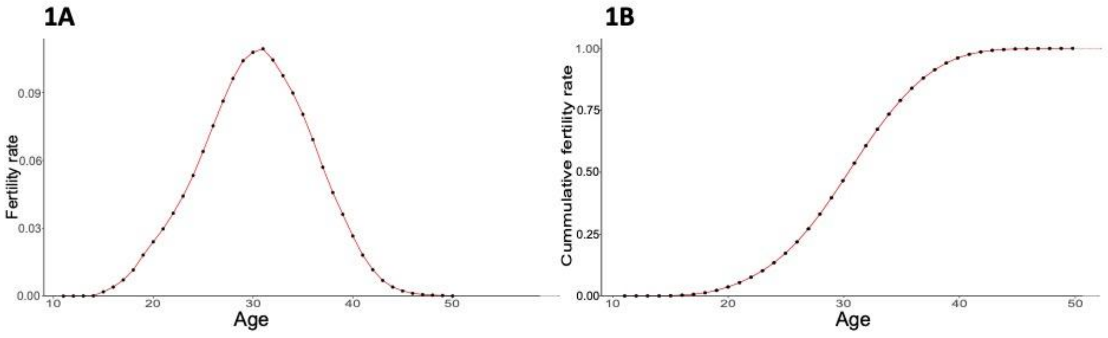


**Figure S2.** Fate of the 1,000 lineages simulated as descendants of one ancestral with a CAG expansion with an initial population frequency of 0.0001. (A) Frequency of the descendant alleles of the original CAG expansion in *ATXN2* founder, per generation (B) Frequency of the descendant alleles of the original CAG expansion in *ATXN3* founder, per generation.

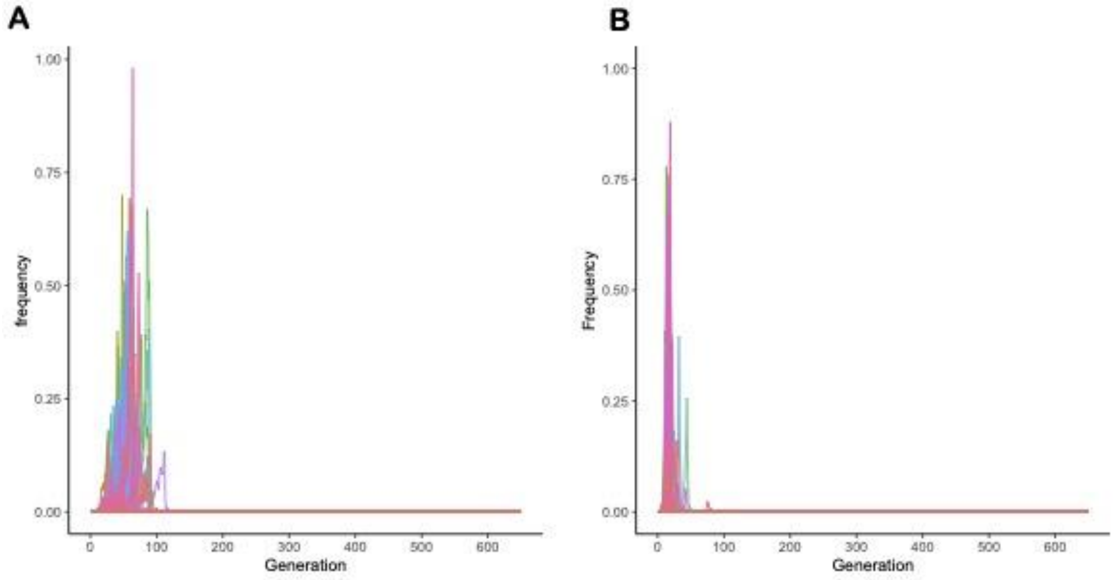

**Figure S3.** The median (IQR) number of generations in which expanded alleles remained present in *ATXN2* and *ATXN3*, in gene dropping simulations.

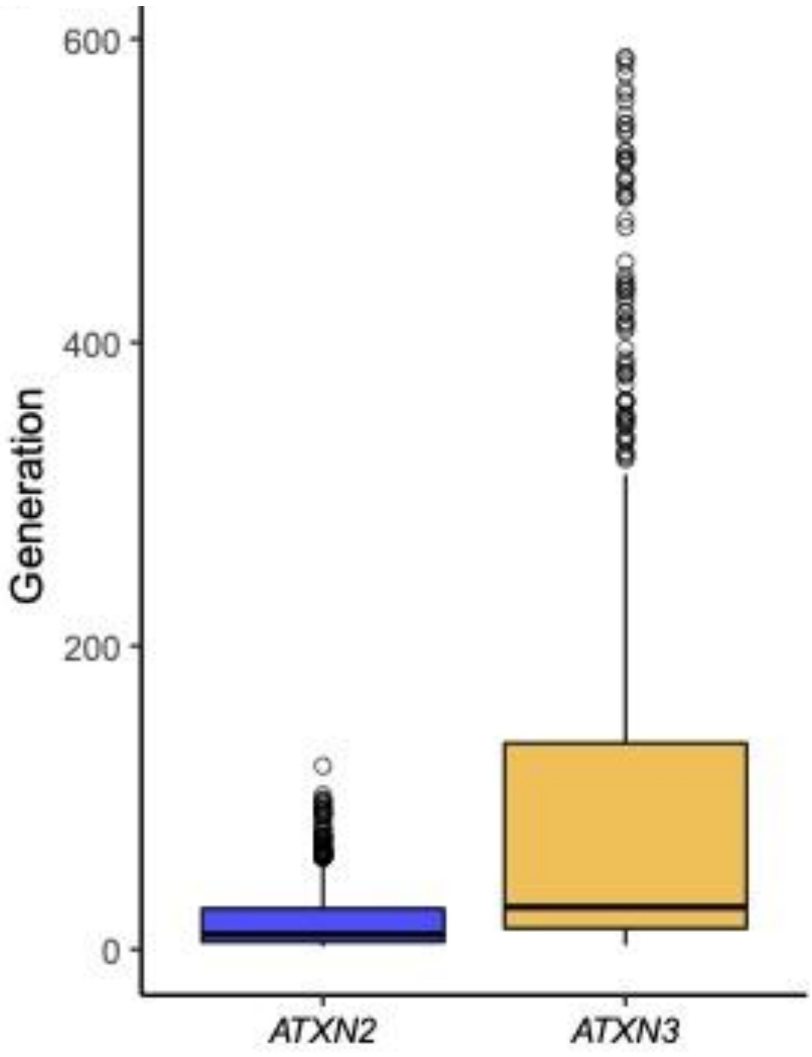
