## Supplementary material for "A model for the dynamics of expanded CAG repeat alleles: *ATXN2* and *ATXN3* as prototypes": Sena et al, submitted (Related Article)

**Supplemental Material - Related Article to Sena et al, "A model for the dynamics of expanded CAG repeat alleles: *ATXN2* and *ATXN3* as prototypes"**

Confidential

**Spinocerebellar ataxia type 2 has multiple ancestral origins. Submitted**

Lucas Schenatto Sena[1,2], Gabriel Vasata Furtado[2], José Luiz Pedroso[3], Orlando Barsottini[3], Mario Cornejo-Olivas[4,5], Paulo Ribeiro Nóbrega[6,7], Pedro Braga Neto[6,8], Danyela Martins Bezerra Soares[8], Fernando Regla Vargas[9,10], Clecio Godeiro[11], Paula Frassinetti Vasconcelos de Medeiros[12], Claudia Camejo[13], Maria Betania Pereira Toralles[14], Nelson Jurandi Rosa Fagundes[1,16], Laura Bannach Jardim[1,2,15,17], Maria Luiza Saraiva-Pereira[1,2,15,18], on behalf of Rede Neurogenetica.

[1] Programa de Pós-Graduação em Genética e Biologia Molecular, Universidade Federal do Rio Grande do Sul, Av. Bento Gonçalves, 9500, 91501-970, Porto Alegre, Brazil

[2] Centros de Pesquisa Clínica e Experimental, Hospital de Clínicas de Porto Alegre, Rua Ramiro Barcelos 2340, 90035-903, Porto Alegre, Brazil

[3] Universidade Federal do Estado de São Paulo, Rua Pedro de Toledo 650, 04039-031, São Paulo, Brazil

[4] Neurogenetics Working Group, Universidad Cientifica del Sur, 19 Panamericana S Avenue, 15067, Lima, Peru, 15067

[5] Neurogenetics Research Center, Instituto Nacional de Ciencias Neuroológicas, 1271 Ancas St, 15003, Lima, Peru,

[6] Setor de Neurologia, Departamento de Medicina Clínica, Faculdade de Medicina, Universidade Federal do Ceará, Rua Professor Costa Mendes, 1608, 60430-140, Fortaleza, CE, Brazil.

[7] Centro universitário Christus, Rua Alexandre Baraúna 949, 60430-160, Fortaleza, CE, Brazil.

[8] Curso de Medicina, Centro de Ciências da Saúde, Universidade Estadual do Ceará, Avenida Dr. Silas Munguba, 1700, 60714-903, Fortaleza, CE, Brazil.

[9] Departamento de Genética e Biologia Molecular, Universidade Federal do Estado do Rio de Janeiro, Rua Frei Caneca 94, 20211-010, Rio de Janeiro, Brazil

[10] Laboratório de Epidemiologia de Malformações Congênitas, Instituto Oswaldo Cruz, Fundação Oswaldo Cruz, Avenida Brasil 4365, 21040-900, Rio de Janeiro, Brazil.

[11] Departamento de Medicina Integrada, Hospital Universitário Onofre Lopes, Avenida Nilo Peçanha, 59012-300, Natal, Brazil

[12] Unidade Acadêmica de Medicina, Hospital Universitário Alcides Carneiro, Universidade Federal de Campina Grande, Rua Carlos Chagas s/n, 58107-670, Campina Grande, Brazil

[13] Facultad de Medicina. Universidad de la República, Avenida General Flores 3461, 11700, Montevideo, Uruguay

[14] Universidade Federal da Bahia, Rua Dr. Augusto Viana, 40110-060, Salvador, Brazil

[15] Serviço de Genética Médica, Hospital de Clínicas de Porto Alegre, Rua Ramiro Barcelos 2340, 90.035-903, Brazil

[16] Departamento de Genética, Universidade Federal do Rio Grande do Sul, Av. Bento Gonçalves, 9500, 91501-970, Porto Alegre, Brazil

[17] Departamento de Medicina Interna, Universidade Federal do Rio Grande do Sul, Rua Ramiro Barcelos 2400, 90035-002, Porto Alegre, Brazil

[18] Departamento de Bioquímica, Universidade Federal do Rio Grande do Sul, Rua Ramiro Barcelos 2600, 90035-003, Porto Alegre, Brazil

**Corresponding author:**

Laura Bannach Jardim
DMI FAMED UFRGS, and
Medical Genetics Service
Serviço de Genética Médica,
Hospital de Clinicas de Porto Alegre
Rua Ramiro Barcelos 2350
90035-003 Porto Alegre, Brazil

**Keywords:** Spinocerebellar Ataxia 2, ancestral origin, polymorphic markers, haplotypes

Number of words in the abstract: 241; in the manuscript: 3,963.

Number of tables: 3

Number of figures: 3

Number of supplemental data: 3

### Abstract

Spinocerebellar ataxia type 2 (SCA2) is an adult onset, dominant neurodegenerative disorder due to expansions of a CAG repeat tract at the *ATXN2* gene. A few studies on ancestral haplotypes were performed so far, and the allele C at rs695871 was always found in SCA2 carriers. We aimed to describe SCA2 ancestral haplotypes constructed based on the SNPs rs9300319, rs3809274, rs695871, rs1236900 and rs593226, using the STRs D12S1329, D12S1333, D12S1672 and D12S1332 to determine their genetic variation. Seventy-seven SCA2 families were recruited from Brazil, Peru and Uruguay; 162 chromosomes from the Brazilian general population and the chromosomes with normal repeats from 101 SCA2 carriers were used as 263 controls. Eleven ancestral haplotypes were found in SCA2 families. The most frequent ones were A-G-C-C-C (46.7% of families), G-C-C-C-C (24.6%) and A-C-C-C-C (10.3%), with assigned risks of being associated with disease of $\delta = 0.326$, 0.197 and 0, respectively. Their mean (sd) CAGexp were 41.68 (3.55), 40.42 (4.11) and 45.67 (9.70) ($p = 0.055$, Kruskal-Wallis), while the mean (sd) CAG lengths at normal alleles were 23.85 (3.59), 22.97 (3.93) and 30.81 (4.27) ($p < 0.001$, Kruskal-Wallis test), respectively. The other SCA2 haplotypes were rare: among them, a G-C-G-A-T was found, evidencing a G allele in rs695871. In summary, our work identified eleven distinct SCA2 haplotypes in Brazilian, Uruguayan, and Peruvian families, including an unexpected SCA2 haplotype with a G allele at rs695871. These results suggest that SCA2 has multiple origins in these populations.

### 1. Introduction

The spinocerebellar ataxia type 2 (SCA2) is a neurodegenerative disorder caused by a dominant expansion of a (CAG)n tract located in the *ATXN2* gene that encodes for a protein named ataxin 2 [1]. The upper limit for the length of normal alleles varies from 32 to 33 CAG repeats [2,3]. Although the (CAG)n is polymorphic in normal chromosomes, the $(CAG)_{22}$ allele is the most prevalent by far, reaching 90.1% of the general population [4, 5].

The expanded CAG repeat (CAGexp) and/or the large polyglutamine (polyQ) tract in the coded protein, become toxic and lead to neurodegeneration primarily in the cerebellum and brainstem [6, 7]. The main symptoms are progressive ataxia and dysarthria, slow saccadic eye movements, and peripheral neuropathy [1]: their mean age of onset (AO) is 33.85 years and correlates with the CAGexp length ($r^2 = 0.577$; Sena et al, 2021) [8]. CAGexp transmissions are unstable, with further expansions being the most frequent variation observed in the offspring [9]. As a consequence, anticipation is common and, in extreme cases, symptoms can start in childhood [10].

Anticipation also leads to a reduction in the reproductive period of CAGexp carriers. In fact, SCA2 individuals with early onset of symptoms have less children than those with late onset [11]. In this sense, anticipation would play a selective pressure against the maintenance of SCA2 in different populations.

Few haplotype studies have been carried out in SCA2 patients. Two reports reconstructed ancestral haplotypes exclusively with short tandem repeats (STRs). Based on D12S1672 and D12S1333, two SCA2 origins were determined in Japan [12], while results based only on D12S1672 suggested a common origin in Cuban, English, Indian and Italian SCA2 families [13]. Three studies reconstructed haplotypes using the same single nucleotide polymorphisms (SNPs) rs695871 and rs695872, finding the C-C haplotype in all SCA2 families from Brazil, India, Italy, and Portugal [14, 15, 16]. Although these results seem to contradict the hypothesis of multiple origins, since they proposed one common haplotype for SCA2 families living in distant geographical regions of the world, it is important to emphasize that the use of few markers makes the studies not very effective in detecting multiple origins of any given trait.

Haplotype reconstruction based on more markers and enhanced resolution is clearly needed to confirm or exclude the C-C haplotype as the proposed common ancestor of alleles causing SCA2. To help with this question, our study aimed to build ancestral SCA2 haplotypes using a larger number of markers in a wide sample of families from Brazil, Peru, and Uruguay.

### 2. Material and Methods

#### 2.1 SCA2 carriers and controls

Subjects with a molecular diagnosis of SCA2 and living in Brazil, Peru and Uruguay between 2010 and 2022, were retrieved from the Rede Neurogenetica database [3], stored in protected files at the Hospital de Clinicas de Porto Alegre. Data about their families and their DNA samples were analyzed in this study. In addition, DNA samples from 81 subjects recruited from the general population of Rio Grande do Sul (RS) were obtained from an anonymous biorepository from our laboratory.

Informed consent was obtained from all individual participants. The study was approved by the Institutional Ethics Committee (Comissão de Ética em Pesquisa do Hospital de Clínicas de Porto Alegre) by the numbers CAAE 0324.1.001.000-06 and 02857818.6.0000.5327, GPPG 2006-0384, 2019-0169 and 2019-0254

### 2.2 Polymorphic markers

Five SNPs and four STR were used as polymorphic markers in this study, spanning a genomic region of 640 Kb (**Figure 1**). SNPs were selected if they had been previously used and if their MAF was of at least 0.35. SNP rs695871 has already been used in previous haplotype studies [14, 15, 16]. SNPs rs9300319, rs3809274 and rs593226 were previously used in studies on positive selection of CAG repeats at *ATXN2* [17, 18]. Finally, rs12369009 was selected due to its linkage disequilibrium with the CAG region. The rs695872 SNP was not genotyped since it is fully linked to rs695871 in more than 1200 chromosomes studied. [14, 15, 16]

STRs D12S1329, D12S1333, D12S1672 and D12S11332 were used previously [13,19,14,15,20], and were added here to build extended haplotypes.

### 2.3 Genotyping

DNA was isolated from peripheral blood leukocytes using standard methods. The CAG repeat length (CAGn) analysis was performed by the polymerase chain reaction (PCR) using fluorescent labeled primers flanking the CAG repeat tract at the *ATXN2* gene, followed by capillary electrophoresis into the 3500 Genetic Analyzer (Thermo Fisher Scientific). Results were analyzed through Microsatellite Analysis Software available at www.thermofisher.com.

The rs9300319, rs3809274, rs695871, rs12369009 and rs593226 SNPs genotyping were performed using TaqMan® SNP Genotyping Assays (assays numbers C___1243837_10, C___1243827_10, C___7524190_30, C__30739839_10 and C___2978539_10, respectively) in a final volume of 8 μL containing 2 ng of DNA, according to the assay protocol (Thermo Fisher Scientific). Amplification was performed in the 7500 Fast Real-Time PCR System® equipment

(Thermo Fisher Scientific) as follows: one cycle of 50°C for 2 min, 95°C for 10 min, followed by 40 cycles of 95 °C for 15 s and 60 °C for 1 min.

Multiplex PCR was performed to identify the number of repeats at each STR. Amplifications were performed using 500 mmol/L dNTPs, 2.5 mmol/L $MgCl_2$, 0.5 mmol/L DMSO, and 2 U of AmpliTaq Gold DNA polymerase (Thermo Fisher Scientific) in 1X AmpliTaq Gold buffer (Thermo Fisher Scientific), and 200 ng of genomic DNA. Four sets of fluorescently-labeled primers were used at 0.2 pmol: D12S1672 (5' CAGGTGTGGTAGCACGA 3'; 5' VIC-CTGGAAATTCACATCTGCTT 3'); D12S1333 (5' FAM-TTCAGGTGGTACAGCCGT 3'; 5' CATCAGAAGGCTTCATAGGAAT 3'); D12S1332 (5' GCCAGGTACAGTGGCTC 3'; 5' PET-CTGGGACCACAGGTGTAG 3') and D12S1329 (5' NED-CCTATCCCACCCAGGC 3'; 5' AGTCTGCCCCAGGCAC 3'). The final volume of reaction was 25 uL. PCR cycling parameters were as follows: initial denaturation for 10 minutes at 94°C, followed by 30 cycles of 30 seconds at 94°C, 45 seconds at 60°C, and 90 seconds at 72°C, and a final extension at 72°C for 5 minutes. An aliquot (5 uL) of amplicon was mixed to Hi-Di™ formamide (Thermo Fisher Scientific) and GeneScan™ 500 LIZ (Thermo Fisher Scientific) in a final volume of 10 uL, and capillary electrophoresis was performed in a 3530xL Genetic Analyzer (Thermo Fisher Scientific). Amplicon lengths were estimated using Microsatellite Analysis Software available at www.thermofisher.com.

**2.4 Data analysis**

Each SCA2 family was represented by one haplotype in the estimates of the relative frequencies of lineages. All chromosomes from subjects from the general population of South Brazil plus chromosomes carrying normal *ATXN2* alleles from up to one subject per Brazilian SCA2 sibship comprised the control chromosomes group, used to estimate the haplotypes present in the general Brazilian population. Brazilian SCA2 families were grouped according to their geopolitical regions of residence, as South, Southeast, Northeast, and North Brazil. The remaining SCA2 families were grouped as Peruvian and Uruguayan families.

Frequencies of SNP alleles obtained in SCA2 families and in the control chromosomes were compared using Fisher's exact test with R Statistical Package.

Haplotypes were then inferred by segregation combined with the use of PHASE v2.1.1 (Stephens et al, 2001) [21]. Reconstructed haplotypes with probabilities greater than 0.6 were used for analyses. Linkage disequilibrium analysis between normal and expanded alleles was tested by Fisher's exact test. Evidence for LD was established using $\delta=(Fd-Fc)/(1-Fc)$, where Fd is the carrier frequency and Fc is the frequency of non-carrier chromosomes [22, 23]. When the haplotypic phase was not determined by the algorithm, chromosomes were not included in the analysis.

The haplotypes reconstructed with the five SNPs only (SNPs haplotypes) were considered the markers of SCA2 lineages and called ancestral haplotypes. The haplotypes reconstructed with the five SNPs plus the four STRs were called the extended haplotypes, and were used to deduce the ancestral variants. The characteristics used to deduce that one extended haplotype was probably the ancestral of a given lineage, were (1) the frequency found in the expanded chromosomes, (2) the number of unchanged alleles, and (3) number of families with extended haplotypes just one step away from that putative ancestral haplotype [24].

Associations between the resulting haplotypes and the number of CAG repeats in normal (*ATXN2*normal) and expanded (*ATXN2*exp) chromosomes were done using all the subjects studied.

Phylogenetic networks were performed through the Network 4.6.1.6 software (http://www.fluxus-technology.com) [25], using microsatellite data. A combined reduced median and median-joining calculation was done to reduce reticulation. At first, phylogenetic networks were drawn using all nine molecular markers; later, STR markers were used to build phylogenies for SNP haplotypes, when possible. In all calculations, ε was set to zero, but different weights were attributed to the mutation rate in each marker. The ε value was adjusted for STRs, in an inversely proportional ratio to their variation in the length of the allele repeat [26].

To estimate genetic distances among SCA2 families, pairwise analyses were performed with Arlequin software version 3.5.2.2 (http://anthro.unige.ch/software/arlequin/software), using the sum of square size difference ($R_{ST}$) as a measure of distance; $R_{ST}$ being an analogue of $F_{ST}$ suited for STR haplotype comparisons [27].

### 3. Results

One hundred and twenty individuals from 77 families carrying an expanded CAG repeat in *ATXN2* were included in this study. Sixty seven families were Brazilian, living in Rio Grande do Sul (22 families), São Paulo (29), Rio de Janeiro (5), Ceará (5), Rio Grande do Norte (2), Paraíba (1), Bahia (1), Pará (1), and Acre (1) states. Six Peruvian and four Uruguayan families were also included. Three out of 22 families from Rio Grande do Sul have been described previously (Ramos et al, 2010) [15]. Samples from more than one related subject were obtained in 18 families (multiplex families).

A total of 241 chromosomes with normal *ATXN2* were used as population controls: 79 of them were the normal chromosomes from SCA2 carriers, and 162 were the chromosomes from 81 subjects recruited from the population of Rio Grande do Sul state.

#### 3.1 SNPs and ancestral haplotypes

Allelic frequencies of rs9300319, rs3809274, rs695871, rs12369009 and rs593226 in the 77 expanded *ATXN2* alleles and in the 263 normal *ATXN2* alleles were described in **Table S1.** The SNPs frequencies found in the expanded *ATXN2* alleles (*ATXN2*exp) were significantly different from those found in the normal *ATXN2* alleles (*ATXN2*normal), with the exception of rs3809274 (**Table S1**). The median (range; variance) number of CAG repeats associated with C and G alleles in normal chromosomes were 24.40(14-33; 24.40) and 22.43 (21-33; 3.36); the $(CAG)_{22}$ was found in 42.55% of C and in 89.45% of G alleles.

The 24 haplotypes found in *ATXN2*exp and *ATXN2*normal alleles are shown in **Table 1,** as well as their frequencies. The geographic distribution of the eleven ancestral haplotypes found in SCA2 families is presented in **Figure 2**. Three ancestral haplotypes were frequently found in SCA2 carriers and in controls. The A-G-C-C-C haplotype was present in SCA2 families from all geographic regions, being the most frequent (46.7%) among them. The attributed risk of A-G-C-C-C being associated with SCA2 was $\delta = 0.326$. G-C-C-C-C haplotype was present in controls and in SCA2 families from Brazil, but not from Peru or Uruguay. G-C-C-C-C had an assigned risk of being associated with SCA2 of $\delta = 0.263$. A-C-C-C-C was found in 10.3% of the SCA2 families; its frequency among controls was similar and was unrelated to an increased risk of SCA2. **Table**

**2** presents the distribution of the CAGn in *cis* with these three ancestral haplotypes, in normal and in expanded alleles.

Other six ancestral haplotypes were found in controls and in few SCA2 families each, so that further analyses were not granted. One of them deserves special attention, since it is the first report of the presence of a G allele in rs695871 in SCA2 carriers: the haplotype G-C-G-A-T, found in three individuals from one SCA2 family of mixed ancestry from Southern Brazil (**Figure 2,** in navy blue, and **Table 1**).

Finally, two ancestral haplotypes were found in SCA2 families but not in controls: A-C-C-A-T and A-G-C-C-T (**Table 1**).

#### 3.2 STRs and extended haplotypes

When all SNPs and STRs were used as markers, 187 distinct extended haplotypes were found in cases and controls. Thirty-one extended haplotypes segregated within SCA2 families (**Table S2)**, while 160 were present in the control group only **(Table S3**). SCA2 families and controls shared four extended haplotypes as follows: 17-17-G-C-G-21-A-T-23, 17-19-G-C-C-21-C-C-23, 17-19-G-G-C-21-C-C-23, and 18-12-A-G-C-22-C-C-26. The former three were considered the ancestors of their lineages.

The CAGn at *ATXN2* of the control individuals that shared the four extended haplotypes with SCA2 carriers are presented in **Table 3.** All control subjects carrying the extended haplotypes considered the SCA2 lineage ancestors 17-19-G-C-C-21-C-C-23 and 17-19-G-G-C-21-C-C-23 had 33 CAG repeats, and two out of 16 controls with 17-17-G-C-G-21-A-T-23 had 31 CAG repeats in *cis.* It is also worthy of note to mention that the CAGn variances of control individuals who shared and share not their extended haplotypes with the SCA2 families were of 28.22 and 7.36, respectively (**Table 3**).

The extended haplotypes found in more than one SCA2 family were 26-12-A-G-C-23-C-C-23, 17-19-G-C-C-21-C-C-23, 17-13-A-G-C-23-C-C-25 and 18-16-A-C-C-17-C-C-23, shared by 18, 14,

4 and 4 families, respectively. Of these, 17-19-G-C-C-21-C-C-23 was the only one found in the control group, where it was related to normal large CAG repeats (**Tables S2 and S3)**.

The haplotype trees of the common lineages A-G-C-C-C, G-C-C-C-C and A-C-C-C-C were further detailed in **Figure 3**. A-G-C-C-C was detected in SCA2 families from all geographic regions under study, and presented a genetic diversity of 0.7397 (+/- 0.074) and average gene diversity over loci of 0.491270 (+/- 0.316391). The most conservative interpretation suggested that 26-12-A-G-C-23-C-C-23 was the founder or the older ancestor of the lineage A-G-C-C-C, since this extended haplotype was the most frequent among SCA2 families, and three families might have derived from it with a distance of one mutational step **(Figure 3A).** G-C-C-C-C (**Figure 3B**) has the lowest genetic diversity - 0.5158 (+/- 0.1316) - and average gene diversity over *loci* - 0.405263 (+/- 0.279203). The extended haplotype 17-19-G-C-C-21-C-C-23 was defined as the ancestor; curiously, it was also shared by SCA2 and normal chromosomes. Finally, the lineage A-C-C-C-C **(Figure 3C**) presented the greatest genetic diversity 0.7778 (+/- 0.110) and average gene diversity over *loci*: 0.486111 (+/- 0.344187).

### 4. Discussion

Eleven different ancestral *ATXN2*exp lineages were identified in our South American cohort: A-G-C-C-C (46.7% of *ATXN2*exp families), G-C-C-C-C (24.6%), A-C-C-C-C (10.3%), G-C-C-A-T (5.1%), G-G-C-A-C (2.5%), A-G-C-C-T (2.5%), and A-C-C-A-T, A-G-C-A-T, G-C-C-A-C, G-C-G-A-T, and G-G-C-C-C (each one of the last five combinations present in one or 1.2% of *ATXN2*exp families). A-G-C-C-T and A-C-C-A-T were missing from our control group; if they were derivatives from one of the former haplotypes, such as A-G-C-A-T, remains to be established. The discovery of a Brazilian SCA2 family carrying the G-C-G-A-T haplotype refuted the current hypothesis that expansions would be universally linked to C allele in rs695871. In general terms, these results denoted that the *ATXN2*exp have multiple origins, and may, rarely, occur in the most protected *ATXN2* background linked to the G allele in the rs695871.

Before our study, the available data on the ancestral origins, population frequencies and genetic characteristics of the *ATXN2*norm and *ATXN2*exp alleles formed a heterogeneous and intriguingly contradictory set. On one hand, SCA2 was characterized by very intense anticipations [28,11, 29, 30], suggesting that the lineages would have rapid extinctions [8]. On the other hand, all previous studies that used standardized markers to reconstruct haplotypes found a single ancestral pattern, the C-C at rs695871 and rs695872 [14, 15, 16], in families living worldwide. The G allele in rs695871 was associated to 70.9% of Indian [14] and to 89.45% of the present Brazilian controls carrying the ($CAG_{22}$) allele. Considering that ($CAG_{22}$) is very stable, descendent chromosomes would have a negligible chance of ever expanding and giving rise to a new SCA2 lineage. In contrast, the variable CAGn related to C-C pattern (or simply C at rs695871) - 77.4% of them different from the ($CAG_{22}$) - would allow for subsequent instabilities and for expansions to reach the range associated with the disease. The data so far obtained "have provided the support for a limited pool of 'ancestral' or 'at risk' haplotypes from which SCA2 disease chromosomes are derived" [14].

In spite of these data, a large number of different ancestral haplotypes was always a reasonable alternative for SCA2, in view of the severity of anticipation phenomena known to occur in this disease.

A mathematical model that combines anticipation, fitness and segregation distortion estimated that the median (range) survival of an expanded allele after its *de novo* appearance would be 10 (1 to 121) generations, or 250 (25 to 3,000) years [31]. Given that SCA2 populations were found worldwide, this short estimated survival of lineages led to the proposal that SCA2 lineages should have multiple origins, related or not to haplotypes predisposed to expansions. Descriptions of *de novo* expansion carriers would favor this hypothesis. However, they are quite rare, such as one individual with 22/35 CAG repeats at *ATXN2* with asymptomatic mother and father carrying 22/22 and 22/32 repeats [32].

Our findings provide further evidence in accordance with these expectations, since eleven ancestral lineages were discovered in our South American cohort. At this point it was not possible to clarify the genetic distances and relationships between all lineages, even if some of them presented similitudes - for instance, if A-G-C-C-T could be a derivative of A-G-C-A-T, or vice-versa. Neither our markers nor the population under study allow any speculation in this direction. That said, we will move on to discuss the results of the most informative ancestral haplotypes found in our study.

A-G-C-C-C was the most frequent haplotype found, being present in 36 (46.7%) SCA2 families living in all regions studied. Since A-G-C-C-C was present in 5.3% of control chromosomes, it was related to a relatively high risk for carriers of having an *ATXN2*exp (the δ of 0.467). 26-12-A-G-C-23-C-C-23 was the most frequent extended haplotype; being so common in our cohort, A-G-C-C-C might well be the most common elsewhere**.** A global study including SCA2 families from Europe, Africa and Asia is needed to clarify the A-G-C-C-C origin.

G-C-C-C-C was the second most common ancestral haplotype found in our SCA2 families, with a widespread geographical distribution across Brazil and Peru. Several genetic characteristics made us think that this haplotype might be also associated with an unstable repeat, prone to expansions and consequently to *de novo* mutations originating from an ancestral *ATXN2*normal haplotype.

G-C-C-C-C was only found in Brazilian SCA2 carriers and had the lowest genetic diversity in SCA2 families, suggesting a common and recent origin. Moreover, the mean (SD) CAGn in unexpanded G-C-C-C-C was larger than the length found in the remaining normal chromosomes (**Table 2**); actually, the seven control chromosomes that carried the extended haplotype 17-19-G-C-C-21-C-C-23 had 33 CAG repeats (**Table 3**), a CAG length that in other populations has been related to SCA2 with very late onset [32, 33, 34]. In contrast, SCA2 individuals with this extended haplotype 17-19-G-C-C-21-C-C-23 carried a mean (SD) of 39.53 (3.65) CAGn, while other SCA2 subjects) carried 42.50 (5.18) repeats ($p = 0.0123$). These findings suggest that G-C-C-C-C expanded repeats might have recently arisen from a normal G-C-C-C-C reservoir linked to normal, long *ATXN2* alleles; probably 17-19-G-C-C-21-C-C-23. We cannot be sure of that, since most of our controls originated from South Brazil (**Table 2**). Similar phenomena were seen in spinocerebellar ataxia type 3, also known as Machado-Joseph disease (SCA3/MJD), where the average length of CAGexp was longer in the A-C-A than in the G-G-T haplotypes, and where larger CAGexp were associated with older lineages and smaller CAGexp, with younger lineages [35]. However, there are some facts hard to be connected with a common and recent origin for all G-C-C-C-C SCA2 carriers from our study: the very large geographical distances between families, ten of them living in South, seven in Southeast, one in Northeast and one in North regions of Brazil (**Figure 2**); and the lack of some connecting branches between extended haplotypes of SCA2 carriers in the phylogenetic tree (**Figure 3B**).

Although this hypothesis remains to be better studied, G-C-C-C-C has some characteristics of a haplotype potentially prone to undergo subsequent expansions towards SCA2. The hypothesis of a predisposing haplotype was raised in early studies on C-C haplotype in *ATXN2* [14]. A similar phenomenon was described in the *HTT* gene, where the A1 and A2 haplotypes were associated with longer normal CAGn and with a greater risk of generating *de novo* mutations in Huntington's disease (HD) [36]. Although plausible, this assumption needs to be confirmed by studying a larger number of *ATXN2* controls from different populational origins.

A-C-C-C-C was found in nine SCA2 families, located in South and Southeast Brazil, as well as in Uruguay. There are no extended A-C-C-C-C haplotypes in common between controls and SCA2 families. This might suggest that this lineage entered the local populations recently - for instance, from a migration originating from the Iberic peninsula or from Italy, both being reasonable alternatives based on the history of European occupation of these Latin American regions. The CAGexp length at A-C-C-C-C was larger, though non significant, than other *ATXN2*exp, and we speculate that A-C-C-C-C expansion followed a mechanism distinct from G-C-C-C-C, as A-C-C-C-C was associated with 22.92 ± 3.93 repeats in normal subjects (**Table 2**).

Among the rare SCA2 haplotypes found here, G-C-G-A-T stood out for the unexpected finding of a G allele in rs695871. As mentioned before, previous published data found that a common haplotype ($C^{rs695871}$-$C^{rs695872}$) was present in 100% of Indian, Portuguese, Italian and three Brazilian families also included in this study [14, 15, 16]. Our G-C-G-A-T family greatly increases the variety of origins of this condition, by identifying a lineage not even outlined.

Nevertheless, we are aware that our study has some limitations. The classic marker in rs695872 was not included in the study. However, it is unlikely that the data obtained with this marker would add new information, as the alleles in rs695871 and rs695872 are 100% linked - i.e., C at rs695871 is always linked to C at rs695872. In addition, CAGexp internal interruptions, in turn, could be very informative for the reconstruction of haplotypes, and this was not addressed by our study.

On the other hand, the history of several bottlenecks to which Latin American populations were exposed in the last and recent 500 years prevented us from performing reliable dating of the detected lineages. There was the Amerindian genocide - bigger in Brazil and Uruguay than perhaps in Peru, countries of origin of SCA2 families included in this study. At the same time, the European occupation was carried out in several relevant migratory waves and even depended on the forced migration of a huge number of enslaved Africans. The last major migrations took place between 1825 and 1905, with the arrival of non-Iberian Europeans mainly from Italy, Germany, Poland, and Ukraine. Numerous founder effects have been proposed for geographic isolates on the continent, including that of SCA2 in Holguin, Cuba [37]. It is plausible to imagine that many of the lineages described here come from recent migrations. It is also possible, but very speculative, to propose an Amerindian origin to G-C-G-A-T. If so many ancestral haplotypes occur in South American families, it is easy to to predict the existence of more lineages among other ethnic groups. The heterogeneity of the South American population contributed to confirm the existence of several origins, by including recent Amerindians, European and African ancestries. However, the South American population structure changed significantly in recent centuries. The study on the origins of SCA2 should have a global scale to allow more consistent datation of lineages.

In conclusion, this work identified eleven distinct ancestral haplotypes in Brazilian, Uruguayan, and Peruvian SCA2 families, suggesting multiple origins for SCA2 in these populations. The existence of multiple origins indicates that *de novo* mutations might play an important role in the maintenance of SCA2 in the population. A global, multicentric study is needed to infer the number of origins of SCA2 and their respective dating, as well as to detect other haplotypes associated with expanded *ATXN2*.

### 5. Declaration of interests

The authors declare no competing interests.

### 6. Acknowledgements

The authors are grateful to the individuals who agreed to participate in this study.
This study was supported by Fundo de Incentivo à Pesquisa do Hospital de Clínicas de Porto Alegre (FIPE-HCPA), grants number GPPG 2006-0384, 2019-0169 and 2019-0254; and DNABank Neurogenetics-INCH Lima-Peru trough grant number ASAP/GP2 - MJFF-023323. LSS, MLSP and LBJ were supported by CNPq.

### 7. Authors contribution

L.S.S., G.V.F., M.L.S.P., and L.B.J. contributed to the conception and design of the study; L.S.S., G.V.F., J.L.P., O.B., M.C.O., P.R.N., P.B.N., D.M.S., F.R.V., C.G., P.F.V.M., C.C., M.L.S.P., L.B.J. contributed to the acquisition of data; L.S.S., G.V.F., and L.B.J. analyzed the data; L.S.S. and L.B.J drafted the text; L.S.S. prepared the figures. All authors reviewed the manuscript.

### 8. References


1 Pulst SM. Spinocerebellar Ataxia Type 2. 1998 Oct 23 [Updated 2019 Feb 14]. GeneReviews® [Internet]. Available from: https://www.ncbi.nlm.nih.gov/books/NBK1275/

2 Gardiner SL, Boogaard MW, Trompet S, de Mutsert R, Rosendaal FR, Gussekloo J, Jukema JW, Roos RAC, Aziz NA.(2019). Prevalence of Carriers of Intermediate and Pathological Polyglutamine Disease-Associated Alleles Among Large Population-Based Cohorts. JAMA Neurol.76(6):650-656. doi:10.1001/jamaneurol.2019.0423

3 de Castilhos RM, Furtado GV, Gheno TC, Schaeffer P, Russo A, Barsottini O, Pedroso JL, Salarini DZ, Vargas FR, de Lima MA, Godeiro C, Santana-da-Silva LC, Toralles MB, Santos S, van der Linden H Jr, Wanderley HY, de Medeiros PF, Pereira ET, Ribeiro E, Saraiva-Pereira ML, Jardim LB; Rede Neurogenetica (2014). Spinocerebellar ataxias in Brazil--frequencies and modulating effects of related genes. Cerebellum 13(1):17-28. doi: 10.1007/s12311-013-0510-y.

4 Andrés AM, Lao O, Soldevila M, Calafell F, Bertranpetit J. (2003). Dynamics of CAG repeat loci revealed by the analysis of their variability. Hum Mutat. 21(1): 61-70. doi: 10.1002/humu.10151.

5 Akçimen F, Ross JP, Liao C, Spiegelman D, Dion PA, Rouleau GA. (2021). Expanded CAG Repeats in ATXN1, ATXN2, ATXN3, and HTT in the 1000 Genomes Project. Movement Disorders 36(2), 514–518. https://doi.org/10.1002/mds.28341

6 Adegbuyiroa A, Sedighia F, Pilkington AW,Groovera S, Legleitera J. (2017). Proteins containing expanded polyglutamine tracts and neurodegenerative disease. Biochemistry: 1199–1217. doi:10.1021/acs.biochem.6b00936.

7 Bunting EL, Hamilton J, Tabrizi SJ (2021). Polyglutamine diseases. Curr Opin Neurobiol. 3;72:39-47. doi: 10.1016/j.conb.2021.07.001.

8 Sena LS, Dos Santos Pinheiro J, Hasan A, Saraiva-Pereira ML, Jardim LB. (2021) Spinocerebellar ataxia type 2 from an evolutionary perspective: Systematic review and meta-analysis. Clin Genet 100(3):258-267. doi: 10.1111/cge.13978.

9 Almaguer-Mederos LE, Mesa JML, González-Zaldívar Y, Almaguer-Gotay D, Cuello-Almarales D, Aguilera-Rodríguez R, Falcón NS, Gispert S, Auburger G, Velázquez-Pérez L. (2018). Factors associated with ATXN2 CAG/CAA repeat intergenerational instability in Spinocerebellar ataxia type 2. Clin Genet 94: 346-350. doi: 10.1111/cge.13380.

10 Mao R, Aylsworth AS, Potter N, Wilson WG, Breningstall G, Wick MJ, Babovic-Vuksanovic D, Nance M, Patterson MC, Gomez CM, Snow K. (2002). Childhood-onset ataxia: testing for large CAG-repeats in SCA2 and SCA7. Am J Med Genet 110(4):338-45. doi: 10.1002/ajmg.10467.

11 Sena LS, Castilhos RM, Mattos EP, Furtado GV, Pedroso JL, Barsottini O, de Amorim MMP, Godeiro C, Pereira MLS, Jardim LB. (2019). Selective Forces Related to Spinocerebellar Ataxia Type 2. Cerebellum 18:188–194. doi: 10.1007/s12311-018-0977-7.

12 Mizushima K, Watanabe M, Kondo I, Okamoto K, Shizuka M, Abe K, Aoki M, Shoji M. (1999). Analysis of spinocerebellar ataxia type 2 gene and haplotype analysis: (CCG)1-2 polymorphism and contribution to founder effect. J Med Genet 36(2):112-4.

13 Pang J, Allotey R, Wadia N, Sasaki H, Bindoff L, Chamberlain S. (1999). A common disease haplotype segregating in spinocerebellar ataxia 2 (SCA2) pedigrees of diverse ethnic origin. Eur J Hum Genet 7: 841–845 doi: 10.1038/sj.ejhg.5200372.

14 Choudhry S, Mukerji M, Srivastava AK, Jain S, Brahmachari SK. CAG repeat instability at SCA2 locus: anchoring CAA interruptions and linked single nucleotide polymorphisms (2001). Hum Mol Genet 10: 2437–2446. doi: 10.1093/hmg/10.21.2437.

15 Ramos EM, Martins S, Alonso I, Emmel VE, Saraiva-Pereira ML, Jardim LB, Coutinho P, Sequeiros J, Silveira I. (2010). Common origin of pure and interrupted repeat expansions in spinocerebellar ataxia type 2 (SCA2). Am J Med Genet B Neuropsychiatr Genet. 5; 153B(2):524-531. doi: 10.1002/ajmg.b.31013.

16 Sonakar AK, Shamim U, Srivastava MP, Faruq M, Srivastava AK. (2021). SCA2 in the Indian population: Unified haplotype and variable phenotypic patterns in a large case series. Parkinsonism Relat Disord 89:139-145. doi: 10.1016/j.parkreldis.2021.07.011. Jul 14. PMID: 34298214.

17 Yu F, Sabeti PC, Hardenbol P, Fu Q, Fry B, Lu X, Ghose S, Vega R, Perez A, Pasternak S, Leal SM, Willis TD, Nelson DL, Belmont J, Gibbs RA. (2005) Positive selection of a pre-expansion CAG repeat of the human SCA2 gene. PLoS Genet. Sep;1(3): e41. doi: 10.1371/journal.pgen.0010041.

18 Chen XC, Sun H, Zhang CJ, Zhang Y, Lin KQ, Yu L, Shi L, Tao YF, Huang XQ, Chu JY, Yang ZQ. (2013). Positive selection of CAG repeats of the ATXN2 gene in Chinese ethnic groups. J Genet Genomics 40(10):543-8. doi: 10.1016/j.jgg.2013.08.003.

19 Saleem Q, Choudhry S, Mukerji M, Bashyam L, Padma MV, Chakravarthy A, Maheshwari MC, Jain S, Brahmachari SK. (2000). Molecular analysis of autosomal dominant hereditary ataxias in the Indian population: high frequency of SCA2 and evidence for a common founder mutation. Hum Genet 106(2):179-87. doi:10.1007/s004390051026.

20 Laffita-Mesa JM, Velázquez-Pérez LC, Santos Falcón N, Cruz-Mariño T, González Zaldívar Y, Vázquez Mojena Y, et al. (2012) Unexpanded and intermediate CAG polymorphisms at the SCA2 locus (ATXN2) in the Cuban population: evidence about the origin of expanded SCA2 alleles. . Eur J Hum Genet. 20(1):41-9. doi: 10.1038/ejhg.2011.154.

21 Stephens M, Smith NJ, Donnelly P. A new statistical method for haplotype reconstruction from population data (2001). Am J Hum Genet 68:978–89.

22 Devlin B and Risch N (1995). A comparison of linkage disequilibrium measures for fine-scale mapping. Genomics 29: 311–322. doi: 10.1006/geno.1995.9003

23 Gaspar, C., Lopes-Cendes, I., Hayes, S., Goto, J., Arvidsson, K., Dias, A., et al. (2001). Ancestral origins of the Machado-Joseph disease mutation: a worldwide haplotype study. Am. J. Hum. Genet. 68: 523–528.

24 Martins S, Calafell F, Gaspar C, Wong VC, Silveira I, Nicholson GA, et al. (2007). Asian origin for the worldwide-spread mutational event in Machado-Joseph disease. Arch Neurol 64(10):1502-8. doi: 10.1001/archneur.64.10.1502.

25 Bandelt HJ, Forster P, Rohl A (1999). Median-joining networks for inferring intraspecific phylogenies. Mol Biol Evol 16: 37–48.

26 Martins S, Calafell F, Wong VC, Sequeiros J, Amorim A. (2006). A multistep mutation mechanism drives the evolution of the CAG repeat at MJD/SCA3 locus. Eur J Hum Genet 14(8):932-40. doi: 10.1038/sj.ejhg.5201643.

27 Excoffier L, Lischer HE (2010). Arlequin suite ver 3.5: a new series of programs to perform population genetics analyses under Linux and Windows. Mol Ecol Resour 10:564–7.

28 Filla A, De Michele G, Santoro L, et al (1999). Spinocerebellar ataxia type 2 in southern Italy: a clinical and molecular study of 30 families. J Neurol 246: 467-471.

29 Gambardella A, Annesi G, Bono F, et al (1998). CAG repeat length and clinical features in three Italian families with spinocerebellar ataxia type 2 (SCA2): early impairment of Wisconsin card sorting test and saccade velocity. J Neurol 245: 647-652.

30 Bürk K, Stevanin G, Didierjean O, et al (1997). Clinical and genetic analysis of three German kindreds with autosomal dominant cerebellar ataxia type I linked to the SCA2 locus. J Neurol 244: 256-261.

31 Sena LS, Lemes R, Furtado GV, Saraiva-Pereira ML, Jardim LB. A model for the dynamics of expanded CAG repeat alleles: *ATXN2* and *ATXN3* as prototypes. Submitted.

32 Futamura N, Matsumura R, Fujimoto Y, Horikawa H, Suzumura A, Takayanagi T. (1998) CAG repeat expansions in patients with sporadic cerebellar ataxia. Acta Neurol Scand 98:55-59.

33 Santos N, Aguiar J, Fernandez J et al (1999). Molecular diagnosis of a sample of the Cuban population with spinocerebellar ataxia type 2. Biotecnol Aplic 16: 219–221

34 Fernandez M, McClain ME, Martinez RA et al (2000). Late-onset SCA2: 33 CAG repeats are sufficient to cause disease. Neurology 55: 569–572.

35 Martins S, Coutinho P, Silveira I, Giunti P, Jardim LB, Calafell F, et al. (2008). Cis-acting factors promoting the CAG intergenerational instability in Machado-Joseph disease. Am J Med Genet B Neuropsychiatr Genet 147B(4):439-46. doi: 10.10 02/ajmg.b.30624.

36 Warby SC, Montpetit A, Hayden AR, Carroll JB, Butland SL, Visscher H, et al. (2009). CAG expansion in the Huntington disease gene is associated with specific and targetable predisposing haplogroup. Am J Hum Genet 84(3):351-66. doi: 10.1016/j.a jhg.2009.02.003.

37 Rodríguez-Labrada R, Martins AC, Magaña JJ, Vazquez-Mojena Y, Medrano-Montero J, Fernandez-Ruíz J, Cisneros B, Teive H, McFarland KN, Saraiva-Pereira ML, Cerecedo-Zapata CM, Gomez CM, Ashizawa T, Velázquez-Pérez L, Jardim LB; PanAmerican Hereditary Ataxia Network (2020). Founder Effects of Spinocerebellar Ataxias in the American Continents and the Caribbean. Cerebellum 19(3): 446-458. doi: 10.1007/s12311-020-01109-7.

### 9. Figures

**Figure 1** - Molecular markers used for haplotype reconstruction in this study.

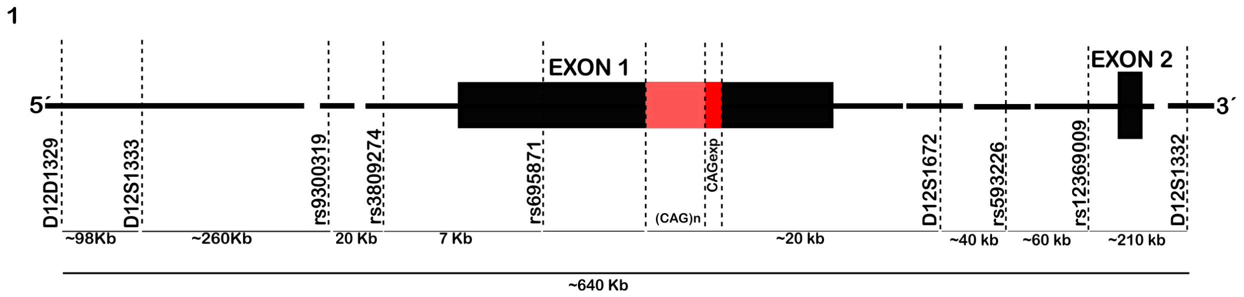

**Figure 2** - Geographic distribution of SNP haplotypes in SCA2 families.

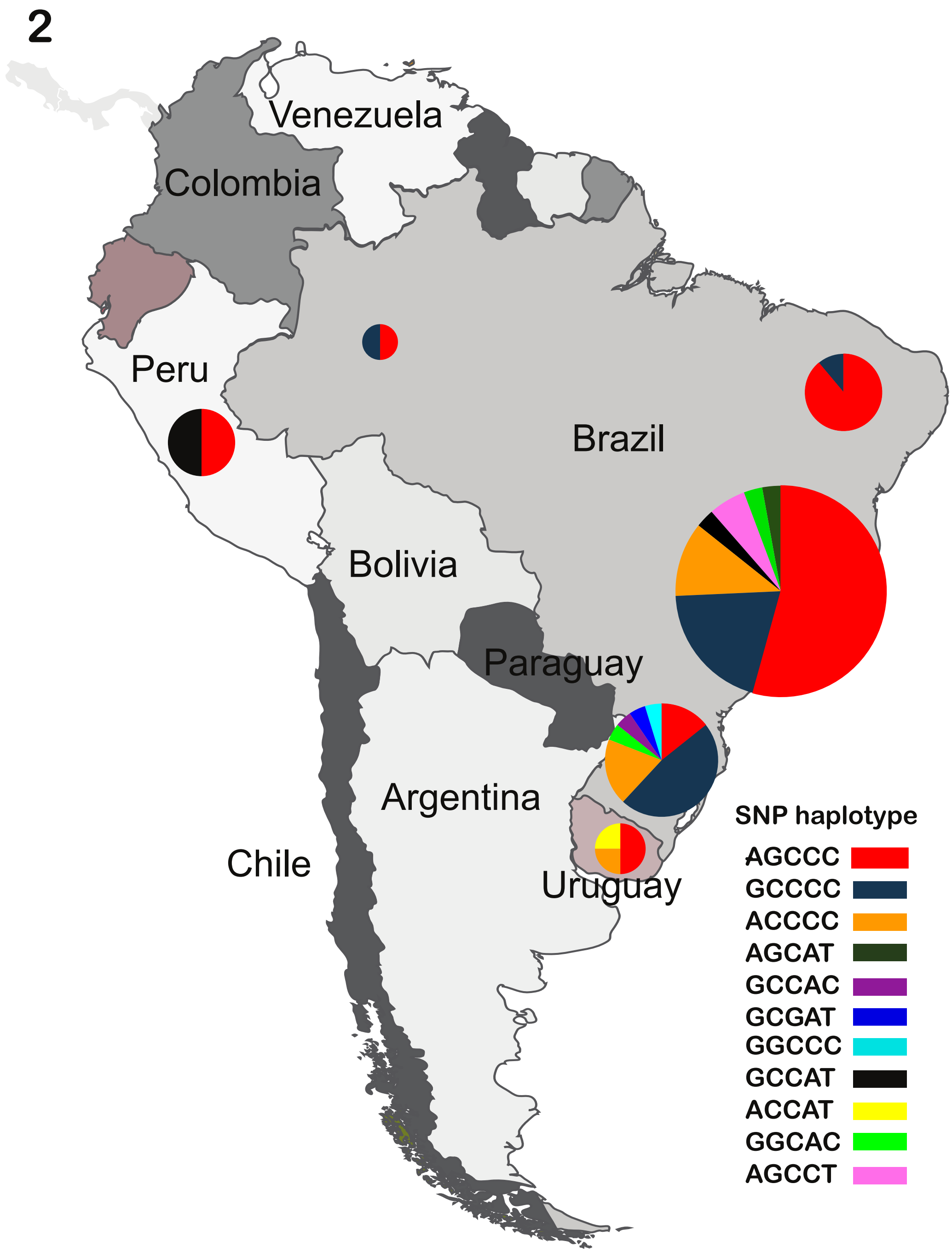

**Figure 3**- Phylogenetic networks of the three haplotypes based on 4 microsatellites. Circles and line sizes are proportional to number of families and stepwise mutation, respectively. The colors in the circles represent the geographical regions where haplotype carriers were living **(A)** A-G-C-C-C haplotype **(B)** G-C-C-C-C haplotype **(C)** A-C-C-C-C haplotype.

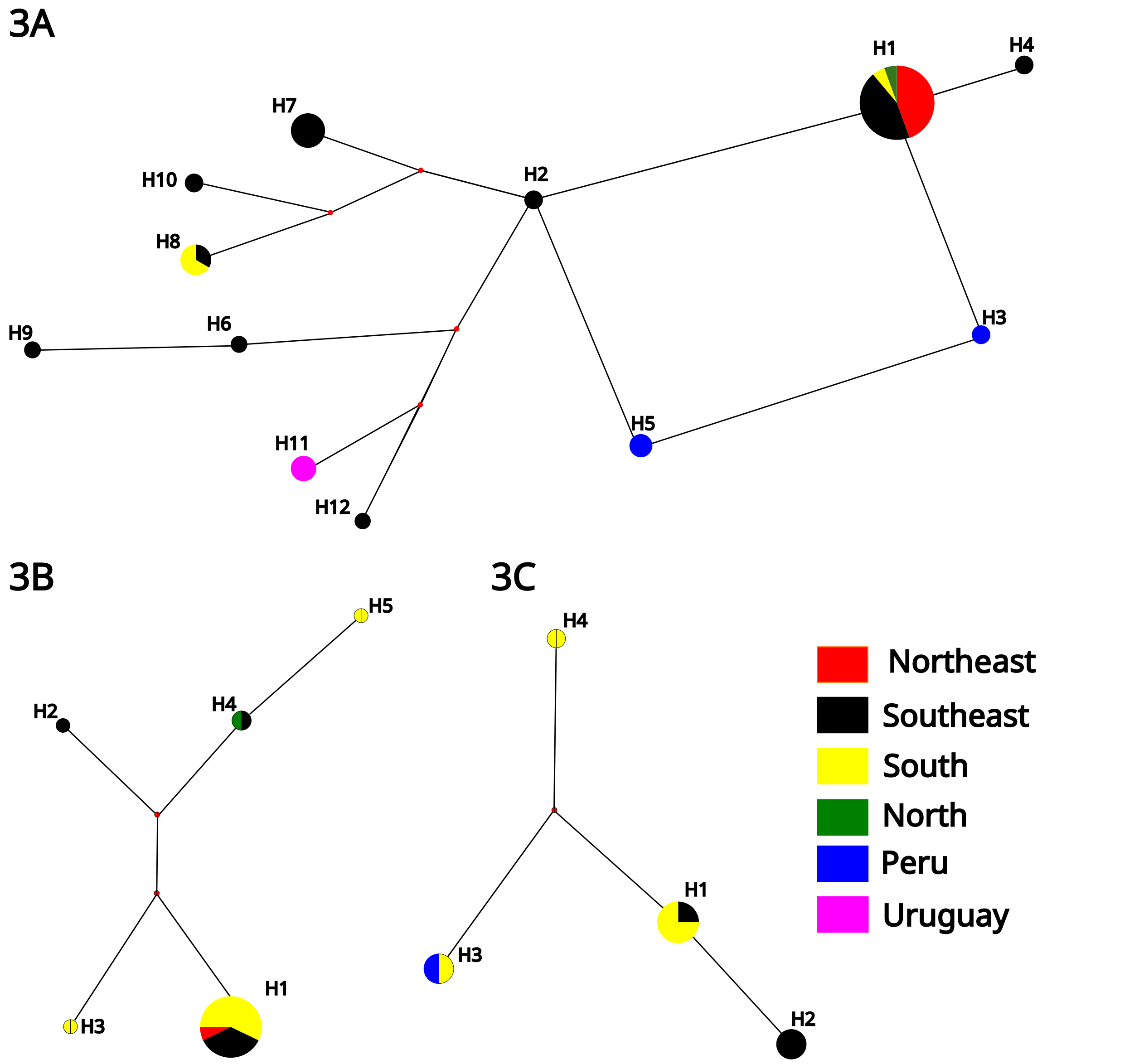

### 10. Tables

**Table 1 - Frequency of SNP haplotypes in SCA2 families and control chromosomes.**

| | *ATXN2exp* (>33 repeats) (total: 77 chromosomes) | | | | *ATXN2* ≤33 (total: 242 chromosomes) | δ* | p** |
|---|---|---|---|---|---|---|---|
| | Number of chromosomes from Brazilian families (total: 67) | Number of chromosomes from Peruvian families (total: 6) | Number of chromosomes from Uruguayan families (total: 4) | Relative frequency (absolute numbers) | Relative frequency (absolute numbers) | | |
| A-G-C-C-C | 31 | 3 | 2 | 0.467(36) | 0.209(55) | 0.326 | <0.001 |
| G-C-C-C-C | 19 | 0 | 0 | 0.246(19) | 0.060(16) | 0.197 | <0.001 |
| A-C-C-C-C | 8 | 0 | 1 | 0.103(9) | 0.053(14) | 0.052 | NS |
| G-C-C-A-T | 1 | 3 | 0 | 0.051(4) | 0.057(15) | - | ns |
| A-G-C-C-T | 2 | 0 | 0 | 0.025 (2) | (0) | 1 | _ |
| G-G-C-A-C | 2 | 0 | 0 | 0.025 (2) | 0.011(3) | 0.014 | ns |
| A-C-C-A-T | 0 | 0 | 1 | 0.012(1) | (0) | 1 | _ |
| A-G-C-A-T | 0 | 0 | 0 | 0.012(1) | 0.007(2) | 0.005 | NS |
| G-C-C-A-C | 1 | 0 | 0 | 0.012 (1) | 0.007(2) | 0.005 | ns |
| G-C-G-A-T | 1 | 0 | 0 | 0.012 (1) | 0.425(112) | _ | _ |
| G-G-C-C-C | 1 | 0 | 0 | 0.012 (1) | 0.022(6) | _ | _ |
| A-C-C-A-C | 0 | 0 | 0 | 0 | 0.007(2) | _ | _ |

| | | | | | | | |
|---|---|---|---|---|---|---|---|
| A-C-G-A-T | 0 | 0 | 0 | 0 | 0.007(2) | _ | _ |
| A-G-C-A-C | 0 | 0 | 0 | 0 | 0.026(7) | _ | _ |
| A-G-G-A-C | 0 | 0 | 0 | 0 | 0.007(2) | _ | _ |
| A-G-G-A-T | 0 | 0 | 0 | 0 | 0.022(6) | _ | _ |
| A-G-G-C-C | 0 | 0 | 0 | 0 | 0.003(1) | _ | _ |
| A-G-G-C-T | 0 | 0 | 0 | 0 | 0.003(1) | _ | _ |
| G-C-G-A-C | 0 | 0 | 0 | 0 | 0.026(7) | _ | _ |
| G-C-G-C-C | 0 | 0 | 0 | 0 | 0.003(1) | _ | _ |
| G-C-G-C-T | 0 | 0 | 0 | 0 | 0.011(3) | _ | _ |
| G-G-G-A-C | 0 | 0 | 0 | 0 | 0.007(2) | _ | _ |
| G-G-G-A-T | 0 | 0 | 0 | 0 | 0.011(3) | _ | _ |
| G-G-G-C-C | 0 | 0 | 0 | 0 | 0.003(1) | _ | _ |

* $\delta$=(Fd-Fc)/(1-Fc) Linkage disequilibrium analysis between expanded(Fd) and normal alleles (Fc)
** Fisher's exact test for linkage disequilibrium.

**Table 2. The mean CAG of the most frequently expanded and normal SNP haplotype**

| | SNP haplotype | n (origin)** | mean CAG(SD) | p * |
|---|---|---|---|---|
| Expanded CAG | A-G-C-C-C | 36 | 41.68 (3.55) | 0.055 |
| | G-C-C-C-C | 19 | 40.42 (4.11) | |
| | A-C-C-C-C | 9 | 45.67 (9.70) | |
| Normal CAG | A-G-C-C-C | 55 (36, 18 and 1 chromosomes from South, Southeast and Northeast Brazil, respectively) | 23.85 (3.59) | < 0.001 |
| | G-C-G-A-T | 112 (83, 17, 4 and 1 chromosomes from South, Southeast, Northeast and North Brazil, and 4 and 3 chromosomes from Peru and Uruguay, respectively) | 22.57 (2.04) | |
| | G-C-C-C-C | 16 (12 and 4 chromosomes from South and Southeast Brazil, respectively) | 30.81 (4.27) | |
| | A-C-C-C-C | 14 (9, 3 and 1 chromosomes from South, Southeast and Northeast Brazil, respectively, and 1 chromosome from Uruguay) | 22.97 (3.93) | |

* Kruskal-Wallis test

** for the control group, only. Please check Table 1 for SCA2 carriers.

**Table 3 - CAG repeat lengths related to the extended haplotypes found in the control group**

| | Haplotype | CAG repeat length mean (variance) | Individual data | |
|---|---|---|---|---|
| | | | CAG repeat lengths found | number of carriers |
| Shared by controls and SCA2 families | 17-17-G-C-G-21-A-T-23 | 23.12(9.45) | 22 | 14 |
| | | | 31 | 2 |
| | 17-19-G-C-C-21-C-C-23 | 33(0.00) | 33 | 7 |
| | 17-19-G-G-C-21-C-C-23 | 33(0.00) | 33 | 2 |
| | 18-12-A-G-C-22-C-C-26 | 22(0.00) | 22 | 1 |
| | all shared haplotypes | 26.5 (28.22) | | |
| Only found in controls | All the others | 22.87(7.36) | | |

### 11. Supplemental Data

**Table S1.** Allelic frequencies of rs9300319, rs3809274, rs695871, rs12369009, and rs593226 in SCA2 families and controls included in this study.

**Table S2.** Extended haplotypes found in SCA2 and in normal chromosomes found in the present study

**Table S3.** All the extended haplotypes found in the present study
